# Relative abundance data can misrepresent heritability of the microbiome

**DOI:** 10.1101/2022.04.26.489345

**Authors:** Marjolein Bruijning, Julien F. Ayroles, Lucas P. Henry, Britt Koskella, Kyle M. Meyer, C. Jessica E. Metcalf

## Abstract

Host genetics can shape microbiome composition, but to what extent it does, remains unclear. Like any other complex trait, this question can be addressed by estimating the heritability (*h^2^*) of the microbiome – the proportion of variance in the abundance of each taxon that is attributable to host genetic variation. However, unlike most complex traits, microbiome heritability is typically based on relative abundance data, where taxon-specific abundances are expressed as the proportion of the total microbial abundance in a sample. We derived an analytical approximation for the heritability that one obtains when using such relative abundances and we uncovered three problems: 1) The interdependency between taxa leads to imprecise heritability estimates. 2) Large sample size leads to high false discovery rates, overestimating the number of heritable taxa. 3) Microbial co-abundances lead to biased heritability estimates. We conclude that caution must be taken when interpreting heritability estimates and comparing values across studies.

## Introduction

The number of host phenotypes known to be impacted by the microbiome is ever-growing, from metabolism to behavior, including its influence on a range of disease risk factors (Cho and Blaser, 2012; Lynch and Hsiao, 2019; Zheng et al., 2020). However, we are only beginning to understand the contribution of host genetics in shaping microbiome composition (Davenport, 2016; Goodrich et al., 2014, 2016; Lopera-Maya et al., 2022; Sanna et al., 2022). This interest stems not only from our desire to understand how evolution and coevolution has shaped host-microbiome interactions over both shorter and longer (i.e. macroevolution) timescales (Dethlefsen et al., 2007; Lynch and Hsiao, 2019), but identifying a genetic basis of host-microbe associations also has important applied health implications (Hall et al., 2017). Critical to address these questions, is our ability to correctly measure the relative importance of hereditary and environmental influences on microbiome composition.

Heritability is a central parameter in quantitative genetics that quantifies a key aspect of the genetic basis for resemblance between parents and offspring. The heritability of a phenotypic trait is a statistical property, defined as the proportion of the phenotypic variance in a population that is attributable to genetic variation (Fisher, 1918). When estimating microbiome heritability, the focal phenotypic trait is typically either a measure of community composition or the abundance of a given taxon (van Opstal and Bordenstein, 2015). While a consensus is emerging that the heritability of most microbiome members is relatively low, specific estimates of the importance of host genetic variation in shaping microbiome composition vary widely across studies (Table 1). For example, a recent study found that 97% of the gut microbes in baboons has a significant non-zero heritability (Grieneisen et al., 2021), while a different study concluded that host genetic background only plays a minor role in shaping microbiome composition in humans (Rothschild et al., 2018). How should we interpret such substantial differences among studies: do these really reflect biological differences?

**Table 1.** Summary of the studies estimating heritabilities of the abundance of microbial taxa, sorted by sample size. More details in Appendix S4 on methodology per study.

| Number | Host system | # Samples | # Taxa | # Heritable taxa | Average non-zero heritability | Host genetic relatedness based on <sup>a</sup> | Normalization / Transformation | Reference |
| --- | --- | --- | --- | --- | --- | --- | --- | --- |
| 1 | Mice | 32 | 43 | NA <sup>b</sup> | 0.47 <sup>b</sup> | Lineage | Total sum scaling | (O'Connor et al., 2014) |
| 2 | Chickens | 56 | 23 | 0 | NA | Pedigree | Log-transformation and scaling | (Zhao et al., 2013) |
| 3<sup>c</sup> | Humans | 93 | 116 | 14 | 0.35 | SNPs | Quantile normalization | (Davenport et al., 2015) |
|  |  | 91 | 104 | 10 | 0.37 |  |  |  |
|  |  | 127 | 102 | 13 | 0.26 |  |  |  |
| 4 | Humans | 108 | 221 | 0 | NA | Twins | Box-Cox transformation | (Goodrich et al., 2014) data from (Turnbaugh et al., 2009) |
| 5 | Humans | 126 | 2,933 | 0 | NA | Twins | Box-Cox transformation | (Goodrich et al., 2014) data from (Yatsunenko et al., 2012) |
| 6<sup>d</sup> | Humans | 244 | 3 | 1 | 0.35 | Twins | Arcsine square root transformation | (Wright et al., 2021) |
|  |  | 88 | 3 | 0 | NA |  |  |  |
| 7 | Humans | 250 | 109 | 11 | NA<sup>e</sup> | Twins | Box-Cox transformation | (Xie et al., 2016) |
| 8 | Humans | 270 | 249 | 26 | 0.58 | Pedigree | Inverse normal transformation | (Turpin et al., 2016) |
| 9 | Switchgrass | 383 | 110 | 21 | 0.24 | SNPs | Total sum scaling | (Sutherland et al., 2021) |
| 10<sup>f</sup> | Cows | 650 | 512 | 39 | NA<sup>e</sup> | SNPs | Quantile normalization | (Wallace et al., 2019) |
|  |  | 200 | 512 | 3 |  |  |  |  |
| 11 | Humans | 485 | 91 | 42 | 0.34 | Twins | Log transformation and scaling | (Gomez et al., 2017) |
| 12 | Humans | 542 | 369 | 85 | 0.27 | Twins | Inverse normal transformation | (Si et al., 2017) |
| 13 | Mice | 592 | 43 | NA<sup>b</sup> | 0.51<sup>b</sup> | SNPs | Total sum scaling | (Org et al., 2015) |
| 14 | Sorghum | 600 | 1189 | 443 | 0.22 | Lineage | Cumulative sum scaling | (Deng et al., 2021) |
| 15 | Humans | 655 | 85 | 52 | 0.24 | Twins | Inverse normal transformation | (Lim et al., 2017) |
| 16 | Humans | 1068 | 21 | 6 | 0.40 | SNPs | Box-Cox transformation | (Ishida et al., 2020) |
| 17 | Humans | 1081 | 909 | 10 | 0.29 | Twins | Box-Cox transformation | (Goodrich et al., 2014) |
| 18 | Humans | 1176 | 209 | 11 | 0.31 | Twins | Inverse rank-sum transformation | (Kurilshikov et al., 2021) |
| 19<sup>g</sup> | Pigs | 1205 | 1678 | 170 | 0.056 | Lineage | Total sum scaling | (Bergamaschi et al., 2020) |
|  |  | 1295 | 1678 | 261 | 0.078 |  |  |  |
|  |  | 1283 | 1678 | 366 | 0.099 |  |  |  |
| 20<sup>h</sup> | Maize | 4866 | 792 | 143 | 0.17 | Lineage | Log transformation | (Walters et al., 2018) |
|  |  | 45 | 2557 | 5 | 0.45 |  |  |  |
| 21 | Humans | 3,261 | 945 | 52 | 0.30 | Twins | Box-Cox transformation | (Goodrich et al., 2016) |
| 22 | Humans | 4,745 | 242 | 31 | 0.20 | Pedigree | Centered log-ratio transformation | (Gacesa et al., 2020) |
| 23 | Baboons | 16,234 | 283 | 273 | 0.068 | Pedigree | Total sum scaling | (Grieneisen et al., 2021) |

<sup>a</sup> Type of host genetic data to estimate heritability. Pedigree: take into account pedigree to estimate narrow-sense $h^2$ . SNPs: incorporate genetic relatedness matrix based on SNPs, to calculate SNP heritability. Lineage: genotype/lineage as random effect, estimates broad-sense $H^2$ . Twins: compare MZ with DZ twins, to estimate broad-sense $H^2$ .
<sup>b</sup> No significance measures are provided. Average heritability is therefore calculated using all estimates.
<sup>c</sup> Analyses are done for winter, summer and both seasons combined
<sup>d</sup> European and African ancestry
<sup>e</sup> Heritability estimates per taxon are not provided.
<sup>f</sup> Two different breeds
<sup>g</sup> Three time points during host development
<sup>h</sup> 2010 and 2015 field study

The use of heritability as a metric is ubiquitous in genetics, yet what it really measures and how it is interpreted remains the source of much confusion. Heritability is by definition a population-specific estimate, and subject to the influence from the environment and the genetic structure of the population. Moreover, detection of a non-zero microbiome heritability does not tell us anything about the mechanisms that cause related individuals to have, on average, more similar microbiomes. Several mechanisms are possible. Microbes might be vertically transmitted from the parents (and typically the mother) to offspring, for example via transfer during vaginal delivery, or via breast milk (Bäckhed et al., 2015; Ferretti et al., 2018); or horizontally transmitted from other family members, perhaps simply due to their proximity (Tung et al., 2015). Both effects could result in tight connections between host and microbial genotypes, and thus inflate heritability. Alternatively, host genotype might directly influence the types of microbes that can establish, as shown in species of woodrats (Weinstein et al., 2021), and this mechanism will also yield high estimates of heritability. Conversely, if heritability is estimated to be zero this need not mean that there is no vertical transmission (or horizontal transmission from relatives), it might simply mean that the effects of the environment are much larger and overwhelm these transmission effects, as has been found in marine sponges (Björk et al., 2019).

A methodological complexity when estimating microbiome heritability, is that the absolute microbial abundances are typically unknown. It is therefore common practice to calculate relative abundances by setting the sum in each sample to 1, generating so-called ‘compositional data’. The inherent problems with compositional data have been acknowledged for some time, and they are known to lead to spurious correlations between variables, even when there exists no correlation at all (Pearson, 1897). This has more recently been discussed in the context of microbial data, for example when testing for differentially abundant microbes across treatment groups (e.g. host disease status) (Carr et al., 2019; Gloor and Reid, 2016; Gloor et al., 2017; Nearing et al., 2021; Rao et al., 2021; Zhou et al., 2021). The estimation of microbiome heritability is rooted in comparison of differential abundances among host genotypes, and could therefore be subject to similar issues. However to date, studies reporting microbe heritability estimates, have not explicitly considered the potential problems associated with the use of compositional data.

We present an approximation of the taxon specific heritability that one obtains when using relative abundances (we call this estimate *φ*^2^). We show that this metric differs from traditional *h^2^* estimates: *φ*^2^ is not simply a function of host genetic and phenotypic variance, but also depends on various other properties of the focal microbe and the rest of the community. Based on this, we identify three main problems that can arise when using relative abundance data to estimate taxon heritabilities. First, as relative abundances inherently covary, a heritable signal for some microbes can lead to spurious heritability estimates of non-heritable microbes or, *vice versa*, non-heritable microbes can mask a genetic signal in heritable microbes. This problem is most apparent for dominant taxa, and the impact of the issue diminishes for low abundance taxa, where the two heritability estimates (*h*^2^ and *φ*^2^) converge. However, a related second problem remains: while the estimated heritability of a non-heritable microbe can become close to zero, it may never completely reach zero. When a large number of host are sampled, even such a very weak (spurious) heritable signal can be highly significant, reflecting greater statistical power. When considering many microbial taxa in a community, the result is a considerable overestimation of the overall proportion of heritable microbes. Third, microbial taxa that covary in abundance (for instance caused by mutualistic or antagonistic interactions), can systematically bias heritability estimates. Depending on the nature and sign of the covariance, this can either mask or inflate true heritability signals. After deriving our approximation for *φ*^2^. we detail each of these problems. We show that our analytical results match results when we estimate heritability by fitting statistical models to simulated datasets. We then discuss empirical heritability estimates obtained from published studies in the light of our results. In the discussion, we outline some solutions that may partly solve the here described issues. We conclude that caution must be taken when interpreting heritability estimates based on relative abundances and comparing values across studies, and that approximations of microbial absolute abundances may help remedy this issue.

### The heritability of a taxon’s abundance

When estimating the heritability of a taxon, one relies on a quantitative genetic model, considering a taxon’s abundance as a quantitative phenotypic trait of the host. The absolute abundance of taxon *i* in host *j* (*P_ij_*) can be written as:

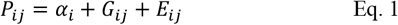

where *α_i_* is the average absolute abundance of microbe *i*, *G_ij_* is the breeding value or host genetic contribution (for simplicity, we assume no genetic dominance or epistasis), *E_ij_* is the environmental contribution (residual), and we assume no G×E interactions. Eq. 1 can be extended by including additional factors that affect taxon abundance, such as host age, sex or season.

Across host individuals, the absolute abundance of microbe *i* is assumed to follow a normal distribution with mean *α_i_* and variance *V_P_i__*. This variance can be decomposed into a genetic and environmental contribution (assuming no genotype-environment covariance):

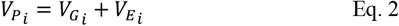

Following the definition of the heritability, the heritability of taxon *i* is:

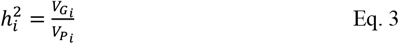

When the absolute abundances are known, one can simply estimate the taxon heritability by quantifying the proportion of the total variance that is attributable to host genetic variation (e.g. by fitting a mixed effects model (Wilson et al., 2010)) (note that in this case, as we assume no dominance or epistasis, the broad-sense and narrow-sense heritability are identical). However, we typically do not know the absolute microbial abundances. Instead, most of the time we quantify how the relative abundance of taxon *i* varies across host individuals and estimate the heritability as the proportion of the variance in relative abundance that is attributable to genetic variation. Below we derive an equation for the obtained heritability when one uses relative, and not absolute abundances, based on the underlying model shown in Eqs 1-2.

### An approximation of the heritability based on relative abundances

As outlined above, the absolute abundance of microbe taxon *i* is distributed across host individuals as:

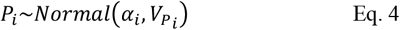

The distribution of relative abundances not only depends on the focal microbe, but also on the absolute abundance of the entire community, consisting of *M* taxa. The community absolute abundance *C* (where 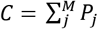) is also a normally distributed variable, where its mean equals the sum of the average abundances over all *M* taxa. The variance depends on the variance in each taxon, plus the sum of each phenotypic covariance between microbial pair, so that:

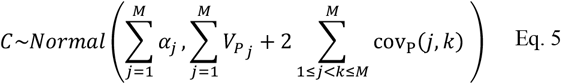

The relative abundance of focal microbe *i* (which we call fraction *f_P_i__*) is calculated as the absolute abundance of focal taxon *i*, divided by the entire community abundance, and therefore is distributed as the ratio between Eq. 4 and Eq. 5:

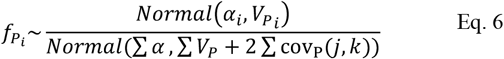

We are interested in quantifying var(*f_P_i__*), as this gives us the total variance in the relative abundance, analogous to *V_P_i__*. Similarly, we can obtain how relative abundances vary between host genotypes, by replacing *V_P_i__* and *V_P_*, by *V_G_i__* and *V_G_*, respectively, and considering genetic covariances covG between each pair of microbes:

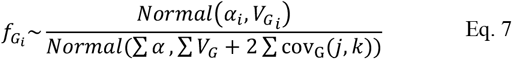

The proportion of the variance in relative abundance explained by host genetic variation (i.e. the heritability based on relative abundances, from now on called *φ*^2^) is then:

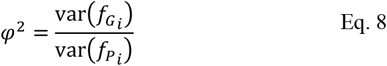

In other words, Eq. 8 gives the heritability that one obtains when using relative, and not absolute, abundances. Ideally, if relative abundances are used as a proxy for absolute abundance, the heritability measure is the same when using absolute and relative abundances, i.e. one hopes that *h*^2^ = *φ*^2^.

An approximation of the heritability of taxon *i* is given by:

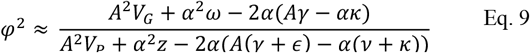

(see Appendix S1). Heritability *φ*^2^ is a function of properties of the focal taxon, with parameters *V_G_* and *V_P_* describing the genetic and phenotypic variance in absolute abundances, and *α* describing the average absolute abundance (to improve readability, we omit subscripts *i*). It follows from Eq. 9 that *φ*^2^ is also a function of the summed genetic and environmental covariances between focal taxon *i* and each of the other taxa in the community (*γ* and *ε*, respectively). Finally, *φ*^2^ is a function of various properties of the background community (excluding the focal taxon): *A* is the average absolute abundance of the background community, *ω* and *z* are the total host genetic and phenotypic variance in absolute abundances of the background community (i.e. the variances summed over all taxa), and *κ* and *ν* are the sums of the genetic and environmental covariances between each pair of background community members. Notice the difference between Eq. 3 and Eq. 9: whereas *h*^2^ is (by definition) only a function of *V_G_* and *V_P_*, the heritability estimate that one obtains when using relative abundances, depends on various additional properties of the focal microbe (*α*), the entire community (*A, ω, z, κ, ν*) and interactions between the focal microbe and the community (*γ*, *ε*). Depending on the biology of the host-microbiome system as well as on properties of the data, we identified three problems that can arise as a consequence.

### Problem 1: Interdependency between taxa leads to imprecise heritability estimates

As relative abundances are not independent, heritable variation in some microbes can lead to spurious non-zero heritabilities, in other microbes. Or *vice versa*, non-heritable microbes can mask a genetic signal in heritable microbes. Consider the extreme scenario with only two equally abundant microbes, where microbe A has a heritability of 1, and microbe B has a heritability of 0 (Fig. 1a). Because abundances are scaled to relative abundances, it would still seem that variation in microbe B abundance is shaped by host genetics (Fig. 1a). Moreover, expressing both abundances as relative abundances partly obscures the host genetic effect on microbe A. This results in a heritability estimate of 0.5 for both species, which is wrong in both cases, and leads to the incorrect conclusion that both microbes are heritable.

**Figure 1.**
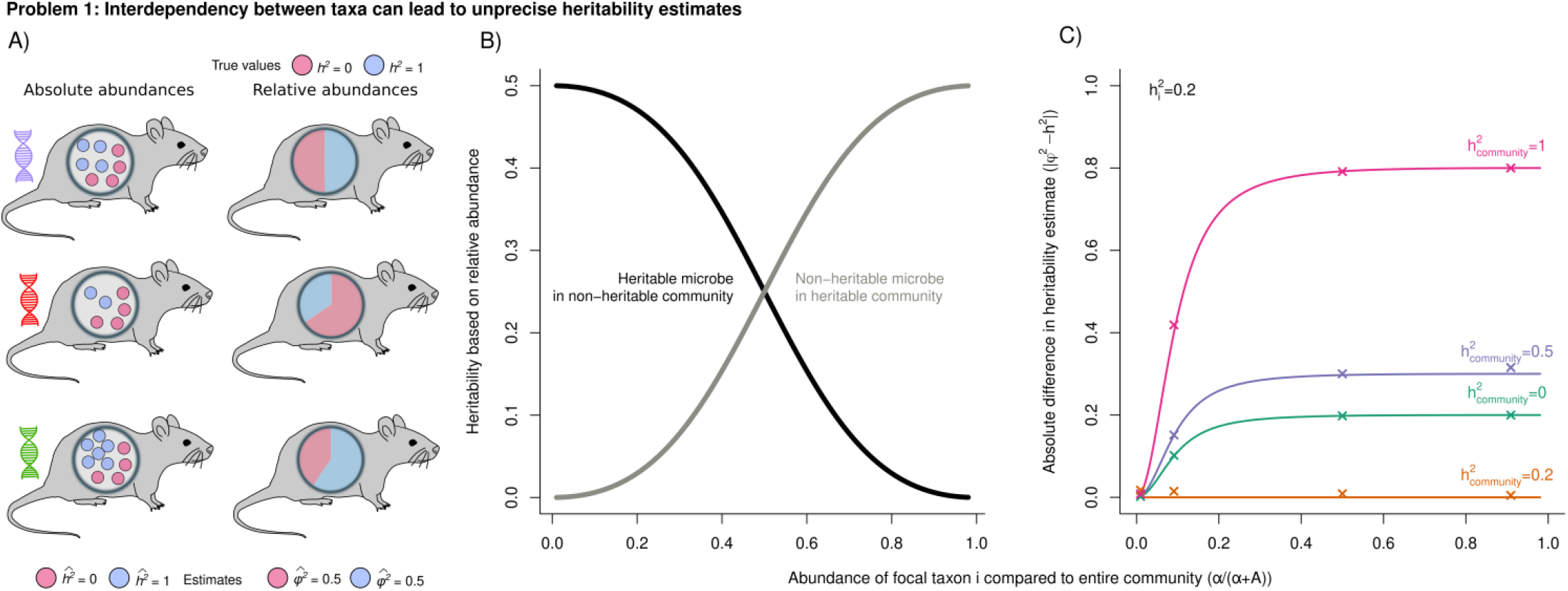
As relative microbial abundances are interdependent, a heritable signal in one microbe can lead to a spurious heritable signal in a second microbe that is not heritable, or mask a genetic signal in a heritable microbe. A) As an example we show three host (mouse) genotypes with two microbes, where one microbe is fully heritable (Blue, h^2^=1), and one microbe is not heritable (Red, h^2^=0). As a consequence, the average absolute abundance of microbe Blue differs among genotypes, while the average abundance of microbe Red is constant. Using the absolute abundances (and with enough host replicates), heritabilities can correctly be estimated. However, as relative abundances are not independent, a host genetic signal in the abundance of the heritable microbe, will also create a host genetic signal in the second microbe, creating variation in relative abundance among genotypes. This leads to an incorrect heritability estimate 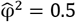 for both microbes. B) When based on relative abundances, properties of both the focal microbe and of the entire community shape the heritability estimates. Here, we vary the average absolute abundance of the focal microbe (α) compared to the absolute abundance of the rest of the community (A) (x-axis shows 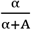). Black line: focal microbe has a heritability of 0.5; the background community is not heritable (A = 1; 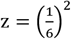; ω = 0; 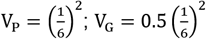). Grey line: focal microbe is not heritable, but the rest of the community has an average heritability of 0.5 (A = 1; 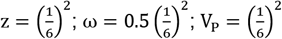; V_G_ = 0). C) Difference in heritability estimates when based on absolute or relative abundances (y axis) when varying *α* compared to A (x axis). When the focal microbe has a low average absolute abundance compared to the total average abundance of the rest of the community (for instance, in the case of many microbial taxa), the difference between *φ*^2^ and *h*^2^ becomes smaller. *h*^2^ of the focal taxon *i* is 0.2, and colored lines show varying heritabilities of the background 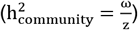. A = 100; 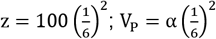. Crosses show results when we estimate heritabilities by fitting a mixed effects model on simulated relative abundance data. To this end, we simulated a population of hosts (500 genotypes × 1000 replicates within each genotype), with microbial communities consisting of 100 taxa (more details in Appendix S2.1-2.3).

This can be formalized using Eq. 9, which, in the absence of genetic and environmental covariances, simplifies to:

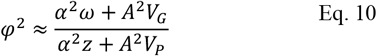

It follows that for a focal taxon with a very low average abundance (i.e. *α* ≪ *A*), the estimated heritability approaches the same value as when based on absolute abundances (Eq. 3):

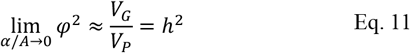

However, for a very dominant taxon (*α* ≫ *A*) it becomes more difficult to retrieve the true heritability *h*^2^, approaching:

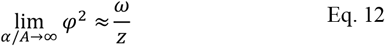

Remember that *ω* and *z* are the total genetic and phenotypic variance of the entire background community (summed over all microbes, excluding the focal microbe). Thus, for a highly dominant microbe, the estimated heritability approaches the heritability of the background community, and is not shaped at all by the genetic and phenotypic variance of the focal microbe.

This implies that depending on properties of both the focal microbe and the rest of the community, heritability estimates can be biased in different directions (Fig. 1b): we will underestimate the heritability of an abundant microbe when it is harbored by a non-heritable community (black line in Fig. 1b). On the other hand, an abundant microbe with no host genetic signal, will still appear heritable when it occurs in the background of a heritable community (grey line in Fig. 1b). As a result, the error in the heritability (i.e. the absolute difference between *φ*^2^ and *h*^2^), depends on both the heritability of the focal microbe, as well as on the heritability of the background community, and in general increases with an increasing abundance relative to the background community (Fig. 1c). When 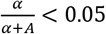 (for instance, in the case of 20 equally abundant taxa in a community), the expected absolute error will be less than 10% for all conditions shown in Fig. 1c. Here we note that the error not only depends on the total abundance of the background community (*A*) compared to the abundance of the focal microbe (*α*), but also on how variances *z* and *V_P_* scale with *A* and *α*, respectively (in Fig. 1c, *V_P_* is kept proportional to *α*).

### Problem 2: Large sample size leads to high false discovery rates

Microbes that are not heritable can still show a genetic signal when abundance measurements are relative, due to the interdependency of the relative abundances. Using Eq. 9 and in the absence of environmental covariances, it follows that the estimated heritability of a non-heritable microbe (by setting *V_G_* = 0) is:

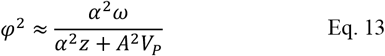

Unless the entire background community is not heritable (i.e. *ω* = 0), Eq. 13 will be larger than 0. Although *φ*^2^ approaches zero when *α* becomes small compared to *A*, it might never reach zero.

Even low *φ*^2^ values can appear significant with enough statistical power. We performed a power analysis using the R-package *simr* (Green and MacLeod, 2016), based on a log likelihood ratio test comparing a model with and without host genetics, to calculate the probability that the null hypothesis (H0: *φ*^2^ = 0) is (wrongly) rejected (Appendix S3 for details). Results again depend on both properties of the focal microbe and of the rest of the community (Fig. 2), but in general, larger sample sizes increase the chance that non-heritable microbes are considered heritable. With a large enough dataset, statistical power reaches 100% (Fig. 2).

**Figure 2.**
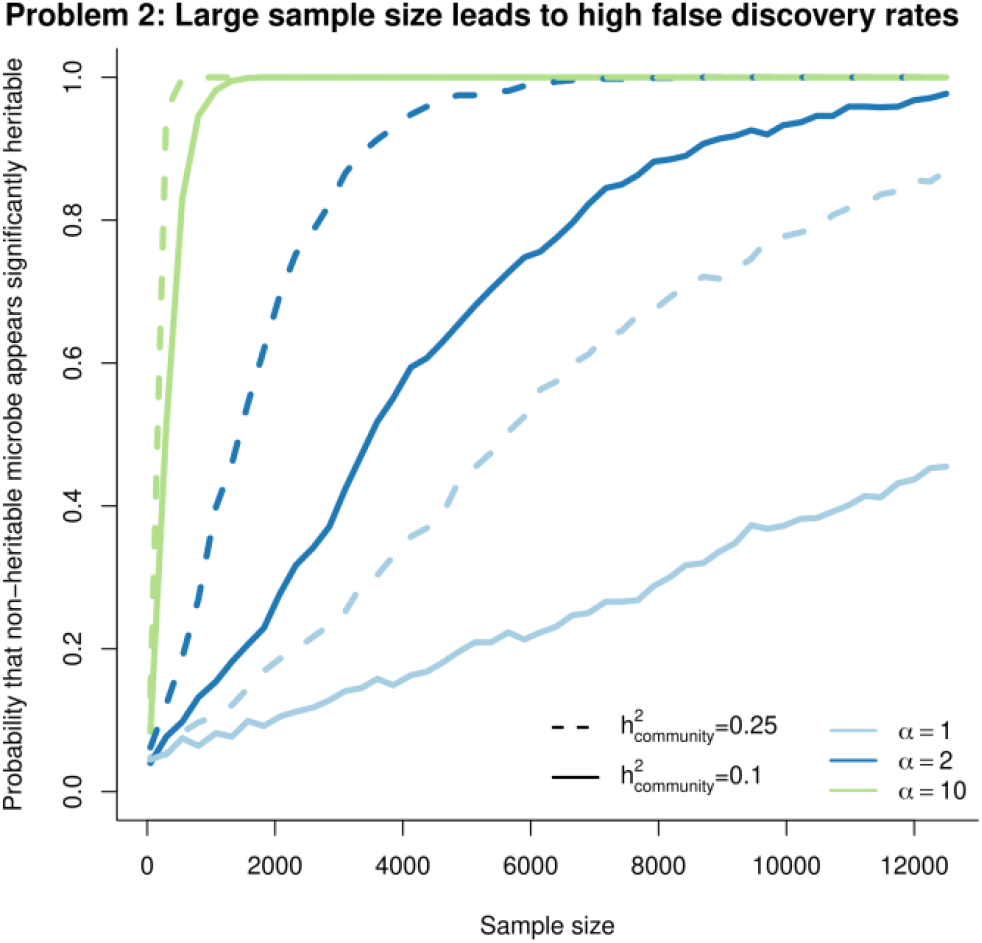
The probability that the heritability of a non-heritable microbe (V_G_ = 0) wrongly appears significant (*α*<0.05) increases with sample size, based on a power analysis using the R-package simr (28). Results depend both on properties of the focal microbe, and on the rest of the community: colors show different abundances of the focal microbe (α) while keeping the background community abundance constant. Line type shows the heritability of the background community (solid lines: 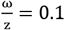; dotted lines: 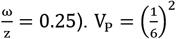; A=100; 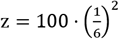.

As a consequence, the number of heritable microbes in a community can be strongly overestimated, especially with a high sample size (more details in results in Appendix S2.4). It is important to note that the high false discovery rates are not a problem of, for instance, sampling error or confounding factors, and increasing data collection efforts or quality alone will not resolve these issues. Similarly, more advanced modeling approaches such as cross-validation, permutation analysis and correcting for multiple testing are unlikely to fully solve this. This is because the problem is inherent to the use of relative abundances: there really *is* a host genetic signal in the relative abundances of non-heritable microbes (i.e. it is not a type 1 error; as Eq. 13 shows, *φ*^2^ really *is* larger than 0).

### Problem 3: Microbial co-abundances lead to biased heritability estimates

Up to this point, we assumed that the covariance terms in Eq. 9 (i.e. *γ*, *ε, ν* and *κ*) were zero. We will now show that relaxing this assumption leads to biased heritability estimates.

Non-zero covariance terms reflect the co-abundance of microbial taxa. In our framing, there are two processes that can cause microbial abundances to covary: host genetic correlations and environmental correlations. The first creates microbial co-abundances at the level of the host genotypes: e.g., a host genotype with an -on average-higher abundance of microbe A, also has a higher abundance of microbe B. The second creates co-abundances at the individual host level, by creating correlated environmental (residual) terms. Note that, as is general practice in quantitative genetics, we use the term ‘environment’ to capture everything outside of genetics: it is essentially a residual term. In the case of the microbiome, it captures not only the effect of ecological environmental factors on microbial abundances, such as temperature or soil, but also effects of the environment inside and shaped by the host, the abundance of other microbes within a host, or simply unexplained noise. One biological process that would lead to the environmental terms being correlated, is microbial interactions. Strong mutualistic interactions, e.g. as a result of cross-feeding or public good production, result in positive environmental correlations. Antagonistic interactions, on the other hand, result in negative environmental correlations.

Non-zero covariances can change heritability estimates in different directions, depending on the nature of the covariance (i.e. genetic or environmental), and whether the covariance involves the focal taxon (*γ, ε*) and/or the background community (*ν, κ*). For the results presented here, we assume that each microbial pair (including focal and background community members) has the same genetic and environmental correlation.

In a community with positive genetic covariances, the heritabilities are generally biased downwards (Fig. 3c). This is because positive genetic covariances have a relatively larger (negative) effect on the numerator than on the denominator (Eq. 9). To make this intuitive, consider the scenario where two equally-abundant microbes both have a heritability of 0.5, and also have a strong genetic correlation (*r_G_*=0.99). Such a strong genetic correlation implies that the host genetic effects for the two microbes covary, so that two microbes show co-abundance at the host genotype level. As a consequence, the absolute abundances vary across host genotypes for both microbes, but they vary in exactly the same way (Fig. 3a). When calculating relative abundances, variation in abundance across genotypes completely disappears, which leads to the incorrect conclusion that none of the microbes show a heritable signal.

**Figure 3.**
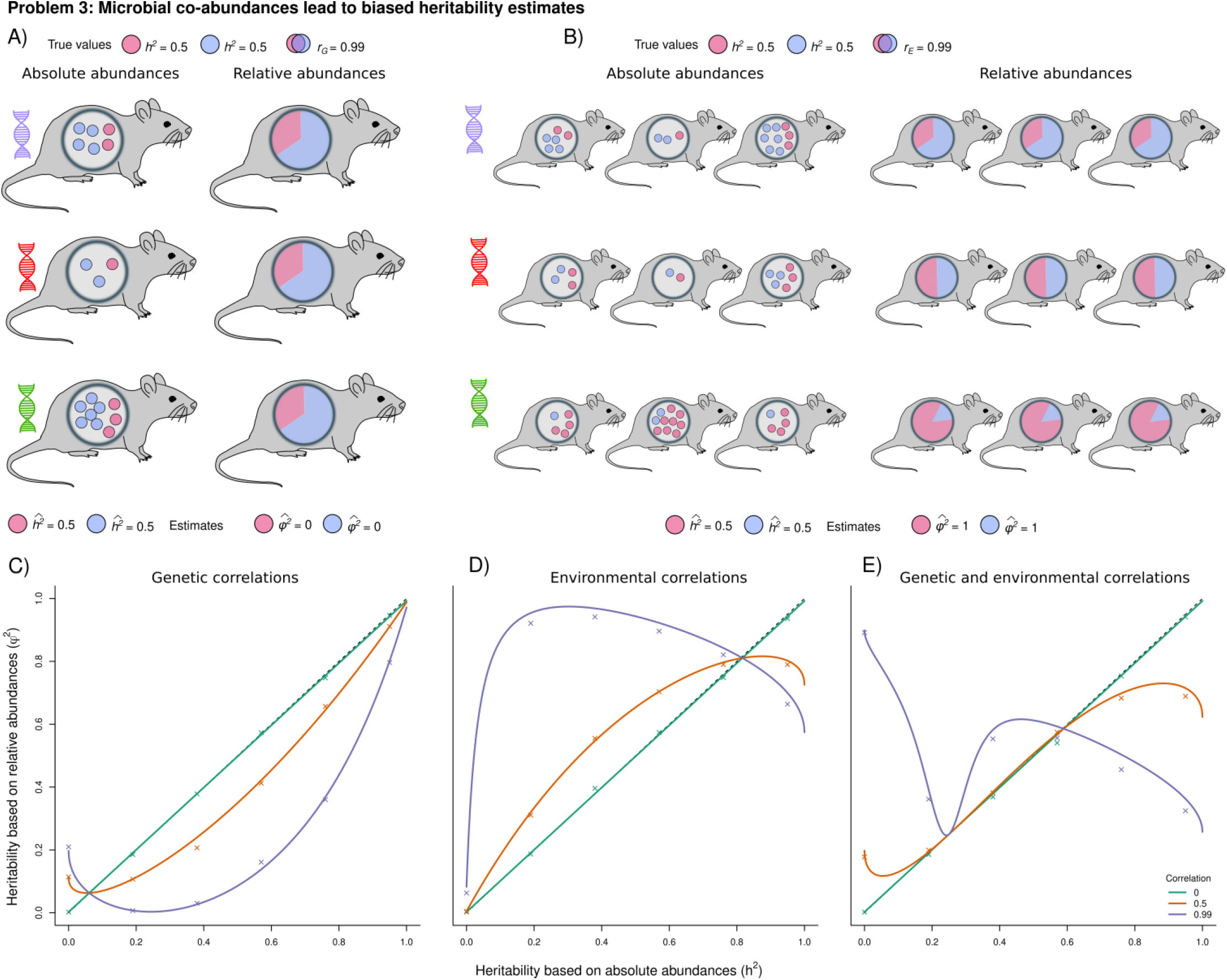
The use of relative abundances leads to biased heritability estimates when there exists host genetic and/or environmental correlations between microbes. A) Illustrates the effects of genetic correlations. As an example we show three host genotypes and two microbes that are both partly heritable (h2=0.5), and with a strong genetic correlation (r_G_=0.99). This implies that host breeding values for the two microbes are strongly correlated. As a consequence, the average absolute abundance of both microbes varies in the same way across host genotypes. Heritabilities can accurately be estimated when using these absolute abundances (estimates for both microbes: 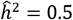). When calculating the relative abundances, however, any variation across host genotypes disappears. This leads to an incorrect heritability estimate 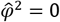 for both microbes, completely masking the host genetic signal. B) Illustrates the effects of environmental correlations. We here show three host genotypes and two microbes that show a strong environmental correlation (r_E_=0.99). As a result, this decreases the amount of variation within genotypes. Heritabilities can be accurately estimated when using the absolute abundances. However, because variation in relative abundance within each genotype is greatly reduced, one obtains a wrong heritability estimate 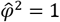 for both species. C-E) Comparison of heritability estimates when based on absolute and relative abundances, varying the environmental correlation (C), the genetic correlation (D) or both (E). *α* = 1; *A* = 100; 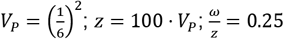. Crosses show results when we estimate heritabilities by fitting a mixed effects model on simulated relative abundance data. To this end, we simulated a population of hosts (500 genotypes ×500 replicates within each genotype) (more details in Appendix S2.5).

The exception is when the true heritability is close to zero: now, positive genetic covariances lead to an overestimation of the true heritability (Fig. 3c). This occurs when *Aγ* < *ακ*, causing the covariance term in Eq. 9 to become negative (thereby increasing the numerator). Since *γ* is the total genetic covariance between taxon *i* and each of the other microbes, it becomes small when *V_G_* is close to zero. As a result, *Aγ* < *ακ*, leading to an overestimated heritability.

Positive environmental covariances in a community (for instance, a highly mutualistic community) has largely opposite effects, by (negatively) affecting the denominator but not the numerator (Eq. 9). Whereas positive host genetic correlations between microbes tend to decrease variation in relative abundance *between genotypes*, positive environmental correlations tend to decrease the amount of variation *within genotypes* (Fig. 3b). When variation within each genotype is reduced, this creates more unique microbiomes to each genotype, suggestive of microbe heritability. As a result, positive environmental covariances lead to a general upward bias in the heritabilities (Fig. 3c). Only if *Aε* < *αν*, the true heritabilities are underestimated. This happens, for instance, if there is little environmental variance in the focal taxon (i.e. a high heritability), causing *ε* to be low.

Finally, when both positive genetic and environmental correlations exist in a community, the relationship between the two heritability measures can become highly non-linear, making it essentially impossible to predict *h*^2^ based on *φ*^2^ (Fig. 3e).

### Framing the current empirical range of estimates

Our results provide additional context in considering the range of estimates of heritabilities published to date. First, our results indicate that estimates of the taxon heritabilities can be precise if each focal taxon has low abundance compared to the total community abundance (and assuming no microbial co-abundances) (Fig. 1c). Our review of the literature indicates that the median number of taxa included in a study is 221 (Table 1). Since most taxa therefore are likely to have low relative abundances, heritability estimates of most individual (low-abundance) taxa may be quite accurate. There is, however, also a wide range in the number of included taxa across studies (varying between 3 and 2933 taxa), and furthermore, human microbiomes are often characterized by a few dominant taxa (Arumugam et al., 2011), and this may be the case for many host species. Our results indicate that for studies that only include a few taxa, or where microbiome communities are characterized by a few highly dominant taxa, precise heritability estimates will be challenging to obtain.

We identified a second problem that is related to the number of sampled hosts: the proportion of heritable microbes can be considerably overestimated due to high false discovery rates. Empirical estimates of the proportion of heritable microbes, show a positive association with the number of hosts sampled (Fig. 4a; binomial regression: p-value < 0.0001). Of course, larger sample sizes always lead to more significant results, as higher sample sizes lead to more power to detect small effects. The challenge here is that without knowing more about the underlying community, we cannot establish how much of this inflation is ‘real’ and how much is due to false discovery. Every microbe may eventually appear significantly heritable with enough statistical power (Fig. 2), even if its absolute abundance is not shaped at all by host genetics. This is due to the interdependency microbiome members will have with other, truly heritable, microbes; and a positive relationship between sample size and the proportion of heritable microbes will emerge even if the true proportion heritable is constant across populations (Appendix S2.4).

**Figure 4.**
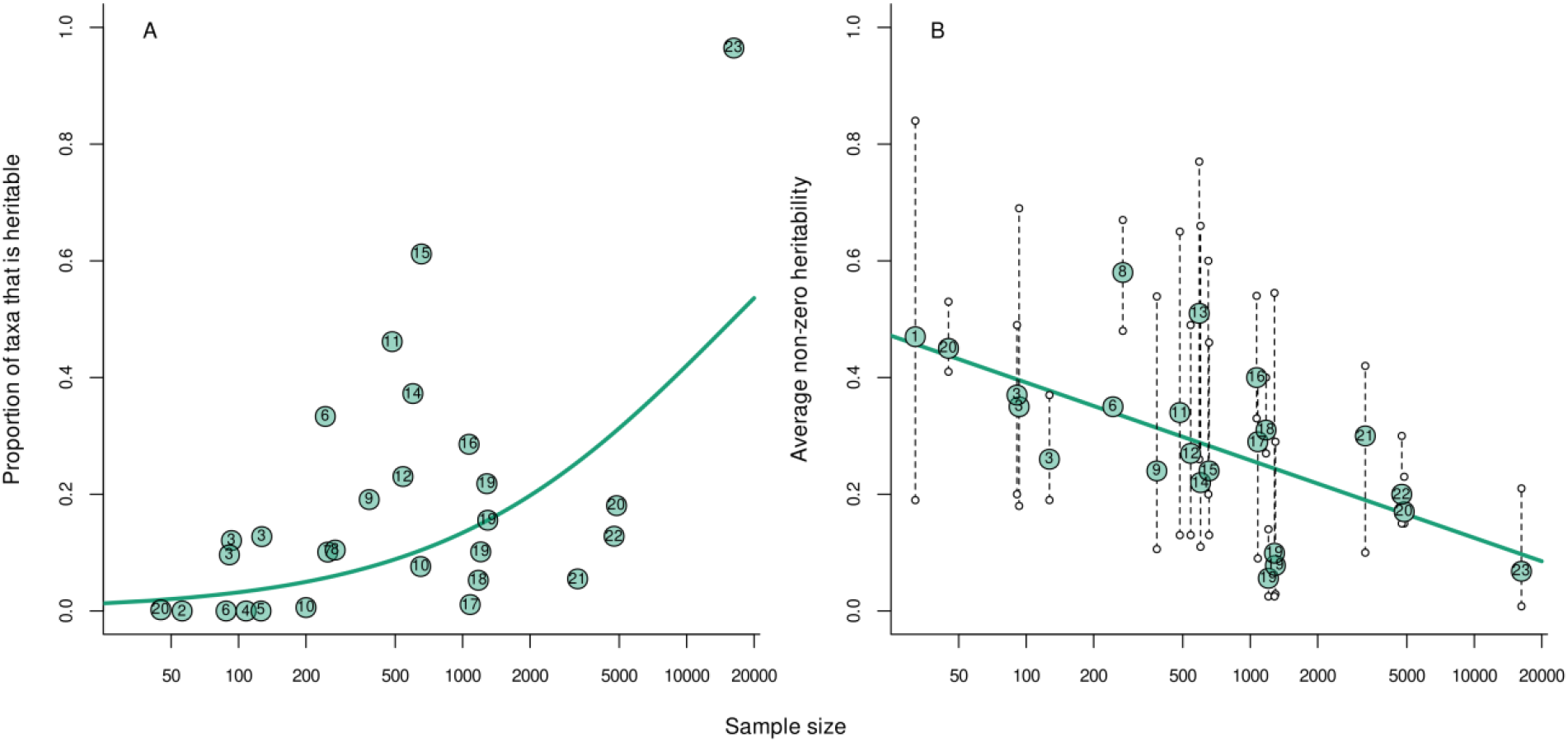
Empirical estimates of the proportion of heritable taxa (A) and the average taxon heritability, including all significantly heritable microbes (B), plotted against sample size, i.e., number of hosts sampled (note the log scale on the x axis). Dots depict values given in Table 1, where the numbers in each dot correspond to the column ‘Number’ in Table 1). Teal lines show the mean prediction based on A) a binomial regression (here the number of trials is the number of taxa), and B) a linear regression. In B) dotted lines connect average heritabilities to the lowest and highest significant heritability found in each study, shown as open dots.

Shifting the focus from the proportion of the taxa that is heritable to considering heritability of taxa, this quantity (including only taxa with a significant heritable signal) varies widely within as well as between studies (Fig. 4b), nearly covering the entire 0-1 range. Across studies, the lowest and highest reported significant heritabilities are 0.008 and 0.84, respectively. The average significant heritability in a community is 0.30, and ranges between 0.056 and 0.58 across studies. It is notable that empirical estimates suggest a negative correlation between sample size and the average heritability, where studies that include a higher number of host individuals report lower average heritabilities (Fig. 4b; linear regression: p-value = 0.002). This could be due to publication bias in smaller studies, in favor of higher heritability estimates, which could suggest that the true microbiome heritabilities may be lower than sometimes reported. However, it could also be that studies with a larger sample size include an increased number of spurious, significant taxa with a low estimated heritability, thereby decreasing the average heritability.

The included studies clearly differ in many aspects other than sample size, both biological (e.g. host system, population and tissue, taxonomic levels, any other covariates) and methodological (e.g. data collection, significance measure, statistical model). There is no reason to expect that the true proportion of heritable microbes or the average heritability is the same across studies - to the contrary. Further, there clearly is variation that is not explained by sample size, indicating that other factors (likely both biological as well as methodological) also play a role. Yet, it is striking that variation in sample size alone, explains considerable variation across studies in both the proportion of heritable microbes (pseudo-R^2^=37%) and in the average heritability (R^2^=39%).

Finally, our results indicate that bias in *φ*^2^ relative to *h*^2^ depends on both the magnitude of *h*^2^ and the underlying pattern of genetic and environmental correlations (Fig 3). Since little is known about the nature and strength of correlations (in absolute abundance) among microbes, it is hard to interpret the impact of this bias on published results to date. Yet, these results do underscore the importance of further efforts to estimate the co-abundance patterns.

## Discussion

Despite the common usage of microbial relative abundance data as a proxy for absolute abundance to estimate microbial heritabilities, few studies have considered the inherent problems that can result from statistical analysis of relative abundances. By their nature, relative abundance data are not independent, creating correlations between variables (microbial abundances) that do not exist in absolute terms. Here we argue that: 1) this can lead to imprecise estimates of heritabilities, especially for microbiomes with fewer taxa and/or highly abundant taxa. 2) Large sample sizes can drive overestimates of the proportion of heritable microbes by increasing the false discovery rate. 3) Patterns of microbial co-abundance, likely to be common in most biological systems, will further bias heritability estimates. Similar challenges have been demonstrated for microbial differential abundance analysis, where it is challenging to control high false discovery rates (Hawinkel et al., 2019; Mandal et al., 2015; Morton et al., 2017; Nearing et al., 2021; Weiss et al., 2016; Zhou et al., 2021). Characterizing the mechanisms underlying these issues helps identify when they might occur, and the direction of bias expected given the number of microbial taxa and their abundance, the number of hosts, and patterns of co-abundance. It is important to note that heritabilities based on relative abundances are potentially misleading only if one wishes to make inferences on host genetic control over *absolute* abundances, i.e., if relative abundances are used as a proxy for absolute abundances. If the metric of interest is, in fact, the heritability of relative abundance, the true value of *φ*^2^ is directly accessible using available relative abundance data. However, as *φ*^2^ is a function of both properties of the focal microbe and of the entire community (Eq. 9), its biological interpretation is potentially challenging. Unfortunately, there does not seem to be a simple solution to fully address the problems described here, but below we discuss several potential approaches for advancing the field.

One solution that would clearly solve the issue of interdependent relative abundance data, is quantifying taxon (or group) absolute abundances. In cases where specific microbial taxa are of interest, such taxa can be directly quantified using such targeted approaches to abundance estimates as quantitative PCR (qPCR), droplet digital PCR (ddPCR), or flow cytometry (Barlow et al., 2020; Rao et al., 2021; Reese et al., 2021; Vandeputte et al., 2017). Additionally, for microbes that are readily cultivable, counts of colony forming units (CFUs) from culturing serve as a method to estimate absolute abundance. However, these approaches remain challenging for microbiome-wide studies that are concerned with the hundreds to thousands of taxa that comprise a given microbiome. One possible solution is to integrate microbial relative abundance data with estimates of the total microbial load of the sample. For instance, if a given taxon represents 1% of the 16S rRNA gene reads in a sample, multiplying that 1% by the total number of 16S rRNA gene amplicons (derived e.g. from qPCR estimates using the same primers, ng of DNA, and PCR cycle numbers), can provide an estimate of that taxon’s absolute abundance. To further improve such an approach, researchers could target known single-copy genes, rather than the 16S rRNA gene, e.g. *rpoB* (Case et al., 2007). Studies that compare inferences when using absolute vs. relative abundances are beginning to emerge (Rao et al., 2021; Vandeputte et al., 2021), although we are not aware of any study that addresses this in the context of microbiome heritability.

In addition to laboratory techniques, new data analysis approaches could prove beneficial. There exists an extensive body of literature on how to analyze compositional data (pioneered by Aitchison [1982]), with relevance to microbiome studies (but also genomics (gene expression), geology (mineral composition) and chemistry (chemical composition)). It is beyond the scope of this paper to provide a comprehensive overview of all available methods, but we refer the interested reader to (Gloor et al., 2017; Hawinkel et al., 2019; Lin and Peddada, 2020a; Quinn et al., 2019) for studies applying such methods to microbial data. Here, we briefly explain the main intuition behind these approaches, and how these may help to improve the accuracy of heritability estimates.

Data normalization is a first solution for obtaining better proxies of the absolute abundances. Instead of dividing the number of reads per taxon by the total number of reads in a sample, one divides the total number of reads by some normalization factor. This involves choosing an appropriate ‘reference’ value, i.e. deciding what the appropriate comparison is within each sample. The advantage of comparing the number of reads for each taxon to a set reference, is that it makes abundances less sensitive to the other taxa that are in the sample. If there are ‘reference’ taxa, known to have constant abundance across samples, one could divide each sample by the number of reads for these reference taxa, thus transforming the relative abundance in each sample into comparable abundances across samples (this is similar to using reference genes to normalize gene expression data). Alternatively, if only a small number of microbial taxa is thought to be differentially abundant across samples, one could also calculate a normalization factor based on some quantile (e.g. median) of each sample’s count distribution (cumulative-sum scaling) (Paulson et al., 2013).

In addition to normalizing, transforming compositional data, e.g., expressing abundances as log ratios is recommended. This transforms data from a simplex to real space, making it more suitable for standard statistical tests (Aitchison, 1982; Greenacre et al., 2021). Different approaches exist, with different reference points: for instance, one could calculate the log-ratio between each taxon and the geometric mean of all taxa (centered log-ratio transformation) (Fernandes et al., 2014), or compare each taxon to a reference taxon (additive log-ratio transformation) (Greenacre et al., 2021; Mandal et al., 2015).

The merit of different normalization and transformation methods critically depends on the chosen reference. If there truly is a known reference taxon with a constant abundance, or if the average abundance truly is identical in all samples, one could successfully correct for sample coverage differences by applying the appropriate normalization/transformation, and retrieve the true heritabilities (Appendix S5). However, while some studies on microbiome heritabilities apply data transformations (e.g. centered log-ratio transformation (Gacesa et al., 2020; Grieneisen et al., 2021), Box-Cox (Goodrich et al., 2014) or inverse normal transformation (Lim et al., 2017)), we lack a validation that such transformations are justified and remedy any existing issues. There is currently little empirical data to guide us in choosing appropriate normalization factors.

It could be more fruitful to focus on the actual heritability estimates, than to focus on the number of significantly heritable taxa. Focusing exclusively on p-values, with some arbitrary threshold for results to be ‘significant’, has been criticized (Halsey et al., 2015; Nakagawa and Cuthill, 2007), and dichotomizing results into ‘significant’ and ‘not significant’ may be particularly problematic for microbiome heritabilities. That is because relative abundances are interdependent: an increase in the abundance of one taxon will inevitably decrease the relative abundance of other taxa. This implies that host genetic variation for the absolute abundance in few microbes, might also lead to genetic variation for other, non-heritable, microbes. Therefore, the null hypothesis (i.e. that there is no host genetic signal in the relative abundances of microbiome members) might rarely be true. With a large enough sample size, this will lead to a statistically significant effect (Nakagawa and Cuthill, 2007) (Fig. 2), even if the effects may be biologically meaningless.

By focusing on effect sizes, we can delineate the heritable taxa that are biologically most relevant. Our results indicate that, unless the focal microbe has a very high abundance compared to the rest of the community (Fig. 1) or microbial abundances covary (Fig. 3), taxon-specific heritability estimates based on relative abundances are unbiased. One could (a priori) set a threshold heritability, and only consider heritabilities exceeding this threshold to be biologically relevant. For instance, Goodrich et al., (2016) only present results of taxa that have an estimated heritability > 0.2.

In addition to focusing on effect sizes, assessing the cumulative evidence for specific microbial taxa will help to identify microbes that are truly heritable and biologically relevant. Grieneisen et al. (2021) found a correlation between their heritability estimates and estimates from previously reported studies (although their effect sizes are much lower). Also, Goodrich et al. (2016) pinpointed various taxa with consistent non-zero heritabilities across studies and across hosts systems. Looking for such consistent results will indicate which taxa merit more detailed study, especially for microbes associated with host performance. Multiple studies have reported high heritabilities for members of the *Christensenellaceae* family, with estimates ranging between 30-60% (Goodrich et al., 2014, 2016; Lim et al., 2017; Turpin et al., 2016; Waters and Ley, 2019). Members of the *Christensenellaceae* have been linked to several host metabolic traits (Waters and Ley, 2019); for example, a higher relative abundance has been associated with a lower body mass index (Goodrich et al., 2014).

In this study, we specifically focused on the consequences of using relative abundances, where the sum in each sample is set to 1, or 100%. The analysis of real-world microbiome datasets comes with additional challenges. First of all, variation across samples not only results in unknown absolute abundances, it also implies different levels of uncertainty. For example, 100 counts of a given taxon in a sample with 10,000 reads, clearly allows for more robust statistical inference than 1 count in a sample with 100 reads, even though the relative abundance in both cases is the same (1%). This information gets lost when converting data into relative abundances.

Further, variation in sampling extent has other important implications. First, we do not know the extent to which we have sampled a host’s microbiome, i.e. what fraction of an individual microbiome was collected for sampling? Knowing the fraction of a microbiome that a sample comprises is crucial to extrapolate absolute abundances to the level of the microbiome (Lin and Peddada, 2020b). Second, we do not know how thoroughly a sample was assessed, i.e. was the number of sequences sufficient to reveal all of a sample’s taxa, or would additional sequencing reveal more taxa? Variation in sampling extent influences the expected number of sampled taxa, where more sequencing reads increases the expected observed microbial richness up to the point of complete assessment (Willis, 2019). Solutions to address this include rarefying (Sanders, 1968), but this is not without criticism (McMurdie and Holmes, 2014). An excess of zero counts results in zero-inflated data, violating the assumption of normally distributed residuals that underlies many parametric statistical tests. Some studies therefore perform log-based transformations to normalize data. However, as we know from community ecology, log transforming count data leads to biased and imprecise estimates, and it involves choosing an arbitrary offset (O’Hara and Kotze, 2010). Further, log-based transformations can lead to incorrect microbiome community-level comparisons, for example resulting in poor estimates of Bray Curtis dissimilarities (McKnight et al., 2019).

How these additional complications further influence the robustness of our microbiome heritability estimates, on top of the issues we describe here, remains to be investigated. With this study, we hope to make researchers aware of the challenges associated with the estimation of microbiome heritabilities. We urge researchers to be careful in interpreting estimates of the heritability of individual taxa, as well as in interpreting the overall proportion of heritable microbes. A focus on consistent results across studies, as well as continued investment in both technical and statistical developments to obtain better approximations of absolute abundances, will likely improve our ability to study the microbiome members that are the most intimately associated with their hosts.

## Acknowledgments

MB is funded by the Dutch Research Council (NWO) (Rubicon grant number 019.192EN.017). BK, KMM and CJEM are funded by the US National Science Foundation (Award number 1754494). LH is funded by the US National Science Foundation (Graduate Research Fellowship Program, #DGE1656466).

## References

Aitchison, J. (1982). The statistical analysis of compositional data. J. R. Stat. Soc. Ser. B 44, 139–160.

Arumugam, M., Raes, J., Pelletier, E., Le Paslier, D., Yamada, T., Mende, D.R., Fernandes, G.R., Tap, J., Bruls, T., Batto, J.-M., et al. (2011). Enterotypes of the human gut microbiome. Nature 473, 174–180.

Bäckhed, F., Roswall, J., Peng, Y., Feng, Q., Jia, H., Kovatcheva-Datchary, P., Li, Y., Xia, Y., Xie, H., Zhong, H., et al. (2015). Dynamics and stabilization of the human gut microbiome during the first year of life. Cell Host \& Microbe 17, 690–703.

Barlow, J.T., Bogatyrev, S.R., and Ismagilov, R.F. (2020). A quantitative sequencing framework for absolute abundance measurements of mucosal and lumenal microbial communities. Nat. Commun. 11, 1–13.

Björk, J.R., Diez-Vives, C., Astudillo-Garcia, C., Archie, E.A., and Montoya, J.M. (2019). Vertical transmission of sponge microbiota is inconsistent and unfaithful. Nat. Ecol. Evol. 3, 1172–1183.

Carr, A., Diener, C., Baliga, N.S., and Gibbons, S.M. (2019). Use and abuse of correlation analyses in microbial ecology. ISME J. 13, 2647–2655.

Case, R.J., Boucher, Y., Dahllof, I., Holmstrom, C., Doolittle, W.F., and Kjelleberg, S. (2007). Use of 16S rRNA and rpoB genes as molecular markers for microbial ecology studies. Appl. Environ. Microbiol. 73, 278–288.

Cho, I., and Blaser, M.J. (2012). The human microbiome: At the interface of health and disease. Nat. Rev. Genet. 13, 260–270.

Davenport, E.R. (2016). Elucidating the role of the host genome in shaping microbiome composition. Gut Microbes 7, 178–184.

Dethlefsen, L., McFall-Ngai, M., and Relman, D.A. (2007). An ecological and evolutionary perspective on human--microbe mutualism and disease. Nature 449, 811–818.

Fernandes, A.D., Reid, J.N., Macklaim, J.M., McMurrough, T.A., Edgell, D.R., and Gloor, G.B. (2014). Unifying the analysis of high-throughput sequencing datasets: characterizing RNA-seq, 16S rRNA gene sequencing and selective growth experiments by compositional data analysis. Microbiome 2, 1–13.

Ferretti, P., Pasolli, E., Tett, A., Asnicar, F., Gorfer, V., Fedi, S., Armanini, F., Truong, D.T., Manara, S., Zolfo, M., et al. (2018). Mother-to-infant microbial transmission from different body sites shapes the developing infant gut microbiome. Cell Host Microbe 24, 133–145.

Fisher, R.A. (1918). The correlation between relatives on the supposition of Mendelian inheritance. Trans. R. Soc. Edinburgh 52, 399–433.

Gacesa, R., Kurilshikov, A., Vila, A.V., Sinha, T., Klaassen, M.A.Y., Bolte, L.A., Andreu-Sánchez, S., Chen, L., Collij, V., Hu, S., et al. (2020). The Dutch Microbiome Project defines factors that shape the healthy gut microbiome. BioRxiv 2020.11.27.401125.

Gloor, G.B., and Reid, G. (2016). Compositional analysis: a valid approach to analyze microbiome high-throughput sequencing data. Can. J. Microbiol. 62, 692–703.

Gloor, G.B., Macklaim, J.M., Pawlowsky-Glahn, V., and Egozcue, J.J. (2017). Microbiome datasets are compositional: and this is not optional. Front. Microbiol. 8, 2224.

Goodrich, J.K., Waters, J.L., Poole, A.C., Sutter, J.L., Koren, O., Blekhman, R., Beaumont, M., Van Treuren, W., Knight, R., Bell, J.T., et al. (2014). Human genetics shape the gut microbiome. Cell 159, 789–799.

Goodrich, J.K., Davenport, E.R., Beaumont, M., Jackson, M.A., Knight, R., Ober, C., Spector, T.D., Bell, J.T., Clark, A.G., and Ley, R.E. (2016). Genetic determinants of the gut microbiome in UK twins. Cell Host Microbe 19, 731–743.

Green, P., and MacLeod, C.J. (2016). simr: an R package for power analysis of generalised linear mixed models by simulation. Methods Ecol. Evol. 7, 493–498.

Greenacre, M., Martinez-Álvaro, M., and Blasco, A. (2021). Compositional Data Analysis of Microbiome and Any-Omics Datasets: A Validation of the Additive Logratio Transformation. Front. Microbiol. 2625.

Grieneisen, L., Dasari, M., Gould, T.J., Björk, J.R., Grenier, J.-C., Yotova, V., Jansen, D., Gottel, N., Gordon, J.B., Learn, N.H., et al. (2021). Gut microbiome heritability is nearly universal but environmentally contingent. Science 373, 181–186.

Hall, A.B., Tolonen, A.C., and Xavier, R.J. (2017). Human genetic variation and the gut microbiome in disease. Nat. Rev. Genet. 2017 1811 18, 690–699.

Halsey, L.G., Curran-Everett, D., Vowler, S.L., and Drummond, G.B. (2015). The fickle P value generates irreproducible results. Nat. Methods 12, 179–185.

Hawinkel, S., Mattiello, F., Bijnens, L., and Thas, O. (2019). A broken promise: microbiome differential abundance methods do not control the false discovery rate. Brief. Bioinform. 20, 210–221.

Lim, M.Y., You, H.J., Yoon, H.S., Kwon, B., Lee, J.Y., Lee, S., Song, Y.-M., Lee, K., Sung, J., and Ko, G. (2017). The effect of heritability and host genetics on the gut microbiota and metabolic syndrome. Gut 66, 1031–1038.

Lin, H., and Peddada, S. Das (2020a). Analysis of microbial compositions: a review of normalization and differential abundance analysis. NPJ Biofilms Microbiomes 6, 1–13.

Lin, H., and Peddada, S. Das (2020b). Analysis of compositions of microbiomes with bias correction. Nat. Commun. 11, 1–11.

Lopera-Maya, E.A., Kurilshikov, A., van der Graaf, A., Hu, S., Andreu-Sánchez, S., Chen, L., Vila, A.V., Gacesa, R., Sinha, T., Collij, V., et al. (2022). Effect of host genetics on the gut microbiome in 7,738 participants of the Dutch Microbiome Project. Nat. Genet. 54, 143–151.

Lynch, J.B., and Hsiao, E.Y. (2019). Microbiomes as sources of emergent host phenotypes. Science 365, 1405–1409.

Mandal, S., Van Treuren, W., White, R.A., Eggesbø, M., Knight, R., and Peddada, S.D. (2015). Analysis of composition of microbiomes: a novel method for studying microbial composition. Microb. Ecol. Health Dis. 26, 27663.

McKnight, D.T., Huerlimann, R., Bower, D.S., Schwarzkopf, L., Alford, R.A., and Zenger, K.R. (2019). Methods for normalizing microbiome data: an ecological perspective. Methods Ecol. Evol. 10, 389–400.

McMurdie, P.J., and Holmes, S. (2014). Waste not, want not: why rarefying microbiome data is inadmissible. PLoS Comput. Biol. 10, e1003531.

Morton, J.T., Sanders, J., Quinn, R.A., McDonald, D., Gonzalez, A., Vázquez-Baeza, Y., Navas-Molina, J.A., Song, S.J., Metcalf, J.L., Hyde, E.R., et al. (2017). Balance trees reveal microbial niche differentiation. MSystems 2, e00162–16.

Nakagawa, S., and Cuthill, I.C. (2007). Effect size, confidence interval and statistical significance: a practical guide for biologists. Biol. Rev. 82, 591–605.

Nearing, J.T., Douglas, G.M., Hayes, M.G., MacDonald, J., Desai, D., Allward, N., Jones, C.M.A., Wright, R., Dhanani, A., Comeau, A.M., et al. (2021). Microbiome differential abundance methods produce disturbingly different results across 38 datasets. BioRxiv.

O’Hara, R., and Kotze, J. (2010). Do not log-transform count data. Nat. Preced. 1.

van Opstal, E.J., and Bordenstein, S.R. (2015). Rethinking heritability of the microbiome. Science 349, 1172–1173.

Paulson, J.N., Stine, O.C., Bravo, H.C., and Pop, M. (2013). Differential abundance analysis for microbial marker-gene surveys. Nat. Methods 10, 1200–1202.

Pearson, K. (1897). Mathematical contributions to the theory of evolution.—on a form of spurious correlation which may arise when indices are used in the measurement of organs. Proc. R. Soc. London 60, 489–498.

Quinn, T.P., Erb, I., Gloor, G., Notredame, C., Richardson, M.F., and Crowley, T.M. (2019). A field guide for the compositional analysis of any-omics data. Gigascience 8, giz107.

Rao, C., Coyte, K.Z., Bainter, W., Geha, R.S., Martin, C.R., and Rakoff-Nahoum, S. (2021). Multi-kingdom ecological drivers of microbiota assembly in preterm infants. Nature 591, 633–638.

Reese, A.T., Phillips, S.R., Owens, L.A., Venable, E.M., Langergraber, K.E., Machanda, Z.P., Mitani, J.C., Muller, M.N., Watts, D.P., Wrangham, R.W., et al. (2021). Age patterning in wild chimpanzee gut microbiota diversity reveals differences from humans in early life. Curr. Biol. 31, 613–620.

Rothschild, D., Weissbrod, O., Barkan, E., Kurilshikov, A., Korem, T., Zeevi, D., Costea, P.I., Godneva, A., Kalka, I.N., Bar, N., et al. (2018). Environment dominates over host genetics in shaping human gut microbiota. Nature 555, 210.

Sanders, H.L. (1968). Marine benthic diversity: a comparative study. Am. Nat. 102, 243–282.

Sanna, S., Kurilshikov, A., van der Graaf, A., Fu, J., and Zhernakova, A. (2022). Challenges and future directions for studying effects of host genetics on the gut microbiome. Nat. Genet. 54, 100–106.

Tung, J., Barreiro, L.B., Burns, M.B., Grenier, J.-C., Lynch, J., Grieneisen, L.E., Altmann, J., Alberts, S.C., Blekhman, R., and Archie, E.A. (2015). Social networks predict gut microbiome composition in wild baboons. Elife 4, e05224.

Turpin, W., Espin-Garcia, O., Xu, W., Silverberg, M.S., Kevans, D., Smith, M.I., Guttman, D.S., Griffiths, A., Panaccione, R., Otley, A., et al. (2016). Association of host genome with intestinal microbial composition in a large healthy cohort. Nat. Genet. 48, 1413–1417.

Vandeputte, D., Kathagen, G., D’hoe, K., Vieira-Silva, S., Valles-Colomer, M., Sabino, J., Wang, J., Tito, R.Y., De Commer, L., Darzi, Y., et al. (2017). Quantitative microbiome profiling links gut community variation to microbial load. Nature 551, 507–511.

Vandeputte, D., De Commer, L., Tito, R.Y., Kathagen, G., Sabino, J., Vermeire, S., Faust, K., and Raes, J. (2021). Temporal variability in quantitative human gut microbiome profiles and implications for clinical research. Nat. Commun. 12, 1–13.

Waters, J.L., and Ley, R.E. (2019). The human gut bacteria Christensenellaceae are widespread, heritable, and associated with health. BMC Biol. 17, 1–11.

Weinstein, S.B., Mart\’\inez-Mota, R., Stapleton, T.E., Klure, D.M., Greenhalgh, R., Orr, T.J., Dale, C., Kohl, K.D., and Dearing, M.D. (2021). Microbiome stability and structure is governed by host phylogeny over diet and geography in woodrats (Neotoma spp.). Proc. Natl. Acad. Sci. 118.

Weiss, S., Van Treuren, W., Lozupone, C., Faust, K., Friedman, J., Deng, Y., Xia, L.C., Xu, Z.Z., Ursell, L., Alm, E.J., et al. (2016). Correlation detection strategies in microbial data sets vary widely in sensitivity and precision. ISME J. 10, 1669–1681.

Willis, A.D. (2019). Rarefaction, alpha diversity, and statistics. Front. Microbiol. 10, 2407.

Wilson, A.J., Réale, D., Clements, M.N., Morrissey, M.M., Postma, E., Walling, C.A., Kruuk, L.E.B., and Nussey, D.H. (2010). An ecologist’s guide to the animal model. J. Anim. Ecol. 79, 13–26.

Zheng, D., Liwinski, T., and Elinav, E. (2020). Interaction between microbiota and immunity in health and disease. Cell Res. 30, 492–506.

Zhou, H., Zhang, X., He, K., and Chen, J. (2021). LinDA: Linear Models for Differential Abundance Analysis of Microbiome Compositional Data. ArXiv Prepr. ArXiv2104.00242.

