## Appendix for "Relative abundance data can misrepresent heritability of the microbiome"

### Supplementary Information for: Relative abundance data can misrepresent heritability of the microbiome

April 25, 2022

**S5 Data transformations** **19**

### S1 Approximating the heritability when based on relative abundances

We here derive an analytical approximation of the heritability that one obtains when using relative abundances, based on a quantitative genetic framing of absolute abundances. According to this framing, the absolute abundance of taxon $i$ in host individual $j$ can be written as:

$$P_{ij} = \alpha_i + G_{ij} + E_{ij} \tag{Eq. S1}$$

$\alpha_i$ is the average absolute abundance of taxon $i$, $G_{ij}$ is a host genetic contribution (i.e. a breeding value; for simplicity, we assume no genetic dominance or epistasis), $E_{ij}$ is an environmental (residual) contribution, and we assume no $G \times E$ interactions.

**Distribution of a taxon's absolute abundance** In a population of hosts, the absolute abundance of each taxon $i$ ($Y$) is distributed as:

$$Y \sim \mathcal{N}(\alpha_i, V_{Pi}) \tag{Eq. S2}$$

where $\alpha_i$ is the average absolute abundance, and $V_{Pi}$ is the total variance, calculated as the sum of the genetic and environmental variance (i.e. $V_{Pi} = V_{Gi} + V_{Ei}$; assuming a zero $G \times E$ covariance).

**Distribution of total community absolute abundance** The absolute abundance of the entire community consisting of $M$ microbes, is also a normally distributed variable which we call $X$. The average of $X$ is simply the sum of the average abundances of each of $M$ microbes: $E[X] = \sum_{j=1}^{M} \alpha_j$. The variance in $X$ depends both on the variance in abundance of each microbe, as well as on the covariance between each pair of microbes:

$$\text{var(X)} = \sum_{j=1}^{M} V_{Pj} + 2 \sum_{1 \le j < k \le M} \text{cov}_\text{P}(j,k) = \sum_{j=1}^{M} V_{Pj} + 2 \sum_{1 \le j < k \le M} [\text{cov}_\text{G}(j,k) + \text{cov}_\text{E}(j,k)] \tag{Eq. S3}$$

Where $\text{cov}_\text{P}(j,k)$ is the phenotypic covariance between taxa $j$ and $k$, which is non-zero when microbial abundances covary. This can also be written as the sum of the genetic and environmental covariances ($\text{cov}_\text{G}$ and $\text{cov}_\text{E}$).

**Distribution of relative abundance** Let us now focus on how the relative abundance of taxon $i$ is distributed across host individuals, based on variables $X$ and $Y$. The relative abundance (fraction $f_{Pi}$) is calculated as the absolute abundance of focal microbe $i$ ($Y$), divided by the entire community abundance ($X$), and therefore is distributed as:

$$f_{Pi} = \frac{Y}{X} \sim \frac{\mathcal{N}(\alpha_i, V_{Pi})}{\mathcal{N}(\sum_{j=1} \alpha_j, \sum_{j=1} V_{Pj} + 2 \sum_{1 \le j < k \le M} \text{cov}_\text{P}(j,k))} \tag{Eq. S4}$$

Eq. S4 gives the total distribution of relative abundances. Similarly, we can also obtain how relative abundances of taxon $i$ vary across host genotypes, by replacing all $V_P$ by $V_G$, and considering the genetic covariances ($\text{cov}_\text{G}$) between each pair of microbes:

$$f_{Gi} \sim \frac{\mathcal{N}(\alpha_i, V_{Gi})}{\mathcal{N}(\sum_{j=1} \alpha_j, \sum_{j=1} V_{Gj} + 2 \sum_{1 \le j < k \le M} \text{cov}_\text{G}(j,k))} \tag{Eq. S5}$$

We are interested in quantifying the variance in $f_{Pi}$ and in $f_{Gi}$, as this is the total, and the genetic variance, respectively, in relative abundances of taxon $i$. The proportion of the variance in relative abundance explained by host genetic variation (i.e. the heritability based on relative abundances, from now on called $\varphi_i^2$) is then:

$$\varphi_i^2 = \frac{\text{var}(f_{Gi})}{\text{var}(f_{Pi})} \quad \text{(Eq. S6)}$$

**Calculating the variance in relative abundance** To calculate var($f_{Gi}$) and var($f_{Pi}$), we use an approximation of the variance of the ratio between two dependent normally distributed variables (Hayya et al., 1975). The variance in $\frac{Y}{X}$, provided that both terms are normally distributed (with mean $\mu$ and variance $\sigma^2$), can be approximated as:

$$\text{var}(\frac{Y}{X}) \approx \frac{\sigma_x^2 \mu_y^2}{\mu_x^4} + \frac{\sigma_y^2}{\mu_x^2} - \frac{2\rho\sigma_x\sigma_y\mu_y}{\mu_x^3} \quad \text{(Eq. S7)}$$

(Hayya et al., 1975). Here, $\rho$ is the correlation between $X$ and $Y$, which we can also write as:

$$\rho = \frac{\text{cov}(X, Y)}{\sigma_x \sigma_y} \quad \text{(Eq. S8)}$$

Since, in our case, we can write $X$ as a function of $Y$ (i.e. $X = Y + \sum_{j \neq i} \mathcal{N}(\alpha_j, V_{Pj})$), the covariance between $X$ and $Y$ is:

$$\text{cov}(X, Y) = \text{var}(Y) + \sum_{j \neq i} \text{cov}_\text{P}(i, j) \quad \text{(Eq. S9)}$$

We now introduce the following 'microbiome community' properties:

$$A = \sum_{\substack{j=1 \\ j \neq i}} \alpha_j \quad \text{(Eq. S10)}$$

$$z = \sum_{\substack{j=1 \\ j \neq i}} V_{Pj} \quad \text{(Eq. S11)}$$

$$\omega = \sum_{\substack{j=1 \\ j \neq i}} V_{Gj} \quad \text{(Eq. S12)}$$

$$\gamma = \sum_{\substack{j=1 \\ j \neq i}} \text{cov}_\text{G}(i, j) \quad \text{(Eq. S13)}$$

$$\epsilon = \sum_{\substack{j=1 \\ j \neq i}} \text{cov}_\text{E}(i, j) \quad \text{(Eq. S14)}$$

$$\kappa = \sum_{1 \leq j < k \leq M} \text{cov}_\text{G}(j, k) \quad \text{(Eq. S15)}$$

$$\nu = \sum_{1 \leq j < k \leq M} \text{cov}_\text{E}(j, k) \quad \text{(Eq. S16)}$$

Eq. S10 - Eq. S16 describe the total average abundance of all non-focal microbes (i.e. excluding taxon $i$) ($A$); the total variance in absolute abundance of all non-focal microbes ($z$); the total genetic variance in absolute abundance of all non-focal microbes ($\omega$); the sum of the genetic covariances between focal microbe $i$ and each of the other community members ($\gamma$); the sum of the environmental covariances between focal microbe $i$ and each of the other community members ($\epsilon$); the sum of the genetic covariances between each pair of background community members ($\kappa$); and the sum of the environmental covariances between each pair of background community members ($\nu$). Doing this has the advantage that we no longer need to consider each of the non-focal microbes explicitly; instead, it is sufficient to know their total effects in terms of abundance and (co)variances. Combining Eq. S7 with the above equations, we can now approximate the total variance in the relative abundance of taxon $i$ as:

$$\mathrm{var}(f_{Pi}) \approx \frac{\alpha^2(z + V_P + 2(\nu + \kappa + \gamma + \epsilon))}{(A+\alpha)^4} + \frac{V_P}{(A+\alpha)^2} - \frac{2\alpha(V_P + \gamma + \epsilon)}{(A+\alpha)^3} \qquad \text{(Eq. S17)}$$

For readability, we have removed subscript $i$ from $\alpha$, $V_G$ and $V_P$. We can rewrite Eq. S17 as:

$$\mathrm{var}(f_{Pi}) \approx \frac{A^2 V_P + \alpha^2 z + 2\alpha^2\kappa + 2\alpha^2\nu - 2\alpha A\gamma - 2\alpha A\epsilon}{(A+\alpha)^4} \qquad \text{(Eq. S18)}$$

Similarly, the variance in relative abundance due to genetic variation can be written as:

$$\mathrm{var}(f_{Gi}) \approx \frac{A^2 V_G + \alpha^2\omega + 2\alpha^2\kappa - 2\alpha A\gamma}{(A+\alpha)^4} \qquad \text{(Eq. S19)}$$

Finall, dividing Eq. S19 by Eq. S18, we obtain the heritability based on relative abundances:

$$\varphi_i^2 \approx \frac{A^2 V_G + \alpha^2\omega + 2\alpha^2\kappa - 2\alpha A\gamma}{A^2 V_P + \alpha^2 z + 2\alpha^2(\kappa + \nu) - 2\alpha A(\gamma + \epsilon)} \qquad \text{(Eq. S20)}$$

### S2 Numerical simulation results

#### S2.1 Microbiome datasets simulation

In order to compare our analytical results to numerical results, we simulated populations of hosts and their microbiomes. For simplicity, we assumed asexual hosts with known genotype identities. Each taxon $i$ out of a total of $M$ taxa can be considered as a taxon from the same taxonomic level (e.g. OTU or genus). We did not simulate taxonomic relationships between microbes, as this is irrelevant for the purpose of our study, however, we note that studies differ in the taxonomic classification levels they consider (Table 1 of the manuscript).

We assigned each taxon an average abundance $\alpha$ and set the standard deviation in absolute abundance proportional to its abundance ($\sqrt{V_P} = \alpha/6$) to ensure that values do not become negative. We assigned each taxon a heritability ($h^2$).

For each taxon $i$ in host individual $k$ that belongs to genotype $j$, we then obtained a microbial (absolute) abundance ($P_{ijk}$), following a standard quantitative genetic framing:

$$P_{ijk} = \alpha_i + G_{ij} + E_{ijk} \tag{Eq. S21}$$

$G_{ij}$ is the genetic effect for microbe $i$ and host $j$ (i.e. a breeding value; varying across genotypes), and $E_{ijk}$ is a residual (environmental) contribution.

Genetic and environmental contributions were sampled from multivariate normal distributions.

$$G_{ij} \sim \mathcal{MVN}(0, \mathbf{K}_G) \tag{Eq. S22}$$

$$E_{ijk} \sim \mathcal{MVN}(0, \mathbf{K}_E) \tag{Eq. S23}$$

Here, $\mathbf{K}_G$ and $\mathbf{K}_E$ are the genetic and environmental variance-covariance matrices, respectively. On the diagonal of these matrices are the genetic and environmental variances ($V_G$ and $V_E$) for each microbe $i$, respectively, calculated as:

$$V_{Gi} = h_i^2 V_{Pi} \tag{Eq. S24}$$

$$V_{Ei} = (1 - h_i^2) V_{Pi} \tag{Eq. S25}$$

To test how genetic and environmental correlations between microbes affect heritability estimates, we set genetic and environmental covariances in abundance on the off-diagonals. The covariance between microbe $i$ and microbe $m$ was calculated as:

$$\mathrm{cov_G}(i, m) = r_G \sqrt{V_{Gi}} \sqrt{V_{Gm}} \tag{Eq. S26}$$

$$\mathrm{cov_E}(i, m) = r_E \sqrt{V_{Ei}} \sqrt{V_{Em}} \tag{Eq. S27}$$

Here, $r_G$ and $r_E$ are the genetic and environmental correlations, respectively, which we varied across simulations.

Finally, per host, we rescaled all abundances to express them as relative abundance by dividing each abundance by the sum of all $M$ microbial abundances in host genotype $j$, individual $k$:

$$f_{ijk} = \frac{P_{ijk}}{\sum_{i=1}^{M} P_{ijk}} \tag{Eq. S28}$$

#### S2.2 Heritability estimation

We used these simulated relative abundances $f$ to fit a linear mixed effects model, per microbial taxon, with host genotype as a random effect: $f_i \sim 1 + (1|Genotype)$. This structure matches fully with the simulation structure; there are no confounding factors; and all abundances are normally distributed. With a sufficiently large data set and when using the true (absolute) abundances as the response variable, we thus retrieve the true coefficients and variance components.

The estimated random effect variance of host genotype measures the variation due to host genetics $V_G$. We calculated heritabilities by dividing $V_G$ by the total variance: $h^2 = V_G/V_P$. Note that as we included only additive genetic effects in our simulations, the broad and narrow sense heritability are identical in our case.

#### S2.3 Problem 1: Interdependency between taxa can lead to unprecise heritability estimates

In Figure 1 of the manuscript, we illustrate how the interdependency between relative abundances can lead to unprecise heritability estimates. For the numerical results in Figure 1c (crosses), we simulated datasets with the following properties. We simulated a microbiome community consisting of 101 taxa (1 focal, and 100 background community members). The average abundance of the focal taxon was either 1, 10, 100 or 1000 (x-axis in Figure 1 of the manuscript), and each community taxon had an average abundance of 1. The focal taxon had a heritability of 0.2, and all background members shared the same heritability (either 0, .2, .5 or 1, see colors in Figure 1 of the manuscript). We simulated microbiome communities for 500 different host genotypes $\times$ 1000 replicated hosts within each population. We calculated heritabilities by fitting a mixed effects model (as explained in Section S2.2).

#### S2.4 Problem 2: Large sample size leads to high false discovery rates

In Figure 2 of the manuscript, we show how larger sample sizes increase the chance that a non-heritable microbe (wrongly) appears significantly heritable. In a community of microbes, this can lead to a great overestimation of the proportion of microbes that is heritable. To illustrate this, we use simulations where we varied the sample size (i.e. number of hosts) and the true proportion of heritable microbes.

When varying sample size, we kept the total number of different host genotypes constant at 50. Having 50 grouping levels is well above the recommended number of levels needed to get reliable random effect variance estimates (Harrison et al., 2018; Oberpriller et al., 2021). Results are thus not driven by error due to limited random effect levels. To vary sample size across simulations, we varied the number of replicated hosts within each genotype between 2 and 1000. This generates datasets on 50 - 50,000 hosts, well covering the range of real-world microbiome datasets used for heritability estimation (Table 1 in the manuscript).

We varied the proportion of microbes that is heritable between 0 and 1, keeping the total number of microbes at 100. For all non-heritable microbes, we set $V_G = 0$. For microbes with a non-zero heritability, we drew values from a uniform distribution, so that for each microbe, $0 < h^2 < 1$.

Per simulated dataset, we then estimated the heritability of each taxon by fitting a mixed effects model as explained in Section S2.2. The significance of the estimated $h^2$ was assessed by comparing the likelihood of this mixed effects model, with that of a simpler model without host genotype. We corrected for multiple testing using the Benjamini Hochberg approach, setting the false discovery rate (FDR) to 0.1. This procedure matches the methods described in a recent study estimating microbiome heritability in baboons (Grieneisen et al., 2021).

The accuracy of identifying the number of heritable microbes was calculated as the F-score, which is the harmonic mean of the precision and recall:

$$\text{Precision} = \frac{\text{TP}}{\text{TP} + \text{FP}} \quad \text{(Eq. S29)}$$

$$\text{Recall} = \frac{\text{TP}}{\text{TP} + \text{FN}} \quad \text{(Eq. S30)}$$

$$\text{F} = 2 \cdot \frac{\text{Precision} \cdot \text{Recall}}{\text{Precision} + \text{Recall}} \quad \text{(Eq. S31)}$$

(TP: True Positives, FP: False Positives, FN: False Negatives). When precision=1, there are no false positives; when recall=1, there are no false negatives. The F-score equals 1 when both precision and recall are 1, implying maximal accuracy.

Under low sample sizes, accuracy in estimating the proportion of heritable microbes is low due to a lack of power to detect significant effects, resulting in type 2 errors (i.e. false negatives), reflected by a recall that is $<1$ (Fig. S1). However, under large sample sizes, accuracy is also strongly reduced (Fig. S1). This is due to a strong reduction in precision (i.e. an increase in false positives), leading to a serious overestimation in the number of heritable microbes (Fig. S1). The reduction in accuracy is strongest when only a small proportion of the microbes is truly heritable; with the largest dataset that we simulated (consisting of 50 genotypes $\times$ 1000 replicated hosts within each genotype = 50,000 samples), the estimated proportion of heritable microbes was about 90%, while in (simulated) reality, only 10% of the microbes was heritable.

When using the (true) absolute abundances, the problem of low statistical power with small data sets (unsurprisingly) remains, but precision does not decrease with sample size, resulting in very accurate estimates of the overall microbiome heritability (Fig. S1). At the core of this issue is the interdependency of relative abundances: a host genetic effect in some microbes, will also create genetic differences in relative (but not absolute) abundances for other, non-heritable microbes. This leads to consistent differences in relative abundance among host genotypes, even if very small, and this is precisely what results in an estimated non-zero genetic variance, appearing significant with enough statistical power (i.e. with enough hosts).

#### S2.5 Problem 3: Microbial co-abundances lead to biased heritability estimates

In Figure 3c-e of the manuscript, we added numerical results (crosses) to our analytical results for a comparison. For these numerical results, we simulated 500 host genotypes and 500 replicated hosts within each genotype. We simulated a microbiome community consisting of 101 microbes (1 focal taxon and 100 background community taxa). We varied both the heritability of the focal taxon (x-axis in Figure 3c-e) and the heritability of each of the background community members (colors in Figure 3c-e). A non-zero genetic correlation creates correlations between breeding values (see Eq. S26), whereas a non-zero environmental correlation creates correlations at the individual host level (Eq. S27). We used the same correlation strength for each pair of microbes to create the covariance matrix.

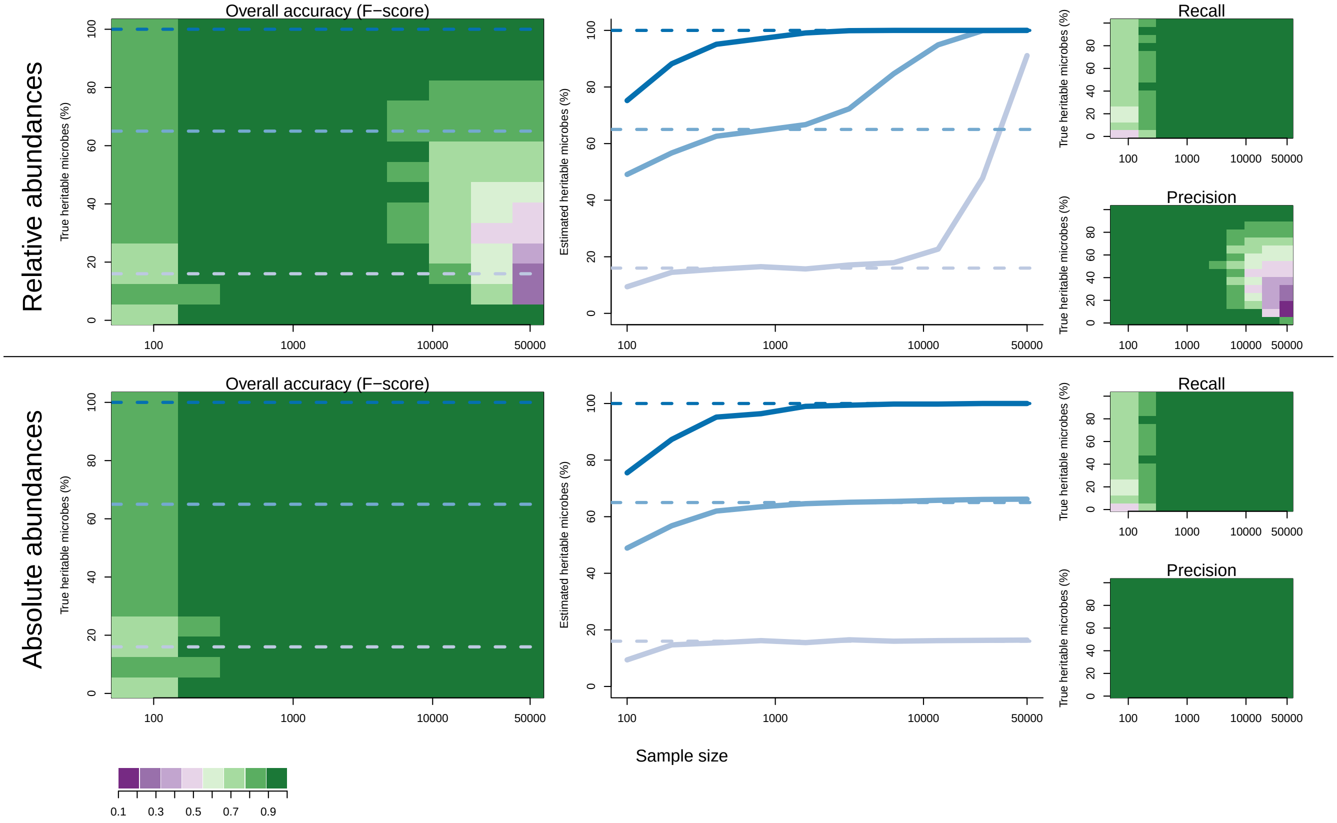

Figure S1: Accuracy (measured as F-score, see Eq. S31) of the estimated percentage of heritable microbes when using relative (upper row) and absolute (bottom row) abundances. The total number of microbes is set to 100 in these simulations. On the left: Accuracy (purple: low; green: high) as a function of sample size (x-axis) and the true percentage of heritable microbes (y-axis). Middle graphs: With small datasets, the true number of heritable microbes (dashed line) is generally underestimated (solid lines below the dashed lines to the left of the plot), due to a lack of statistical power, both when using absolute and relative abundances. Under high sample sizes, the number of heritable microbes is greatly overestimated when using relative abundances, but not when using absolute abundances (solid lines above dashed lines to the right of the plot on the upper panel). On the right: Accuracy split into effects through recall (Eq. S30), and precision (Eq. S29). Recall is the ratio between the number of true positives and the true number of heritable microbes. When recall $< 1$, this indicates the presence of false negatives. Precision is the ratio between the number of true positives and the total number of significant results. A value lower than 1 indicates that there are false positives. Under a high sample size and when using relative abundances, precision is greatly reduced, indicating an increase in false positives. All value are median values of 10 simulations.

#### S3 Power analysis

To calculate the chance that a non-heritable microbe wrongly appears significantly heritable, we performed a power analysis using R-package *simr* (Green and MacLeod, 2016). To this end, we used the function *makeLmer()* to build an artificial *lme4* object. Here, the fixed intercept was the average relative abundance of our focal taxon, calculated as $\frac{\alpha}{\alpha+100}$ (we tested different $\alpha$ values, see colored lines in Figure 3 of the manuscript). We set the slope to 0. The random effect variance ($V_G$) equaled the genetic variance in relative abundance (var($f_{Gi}$); Eq. S19), and the residual variance was calculated as: $\text{var}(f_{Ei}) = \text{var}(f_{Pi}) - \text{var}(f_{Gi})$ (see Eq. S19, Eq. S18). We created a dataset with 25 host genotypes, and $x$ replicates within each host genotype. We used the function $powerCurve()$ to calculate the power at a range of $x$ values (x-axis in Figure 3 of the manuscript shows total sample size, i.e. $x \times 25$). We tested the significance of the random effect by setting the argument *test=random()*, which performs a likelihood ratio test on model with and without genotype as random effect. We performed 1000 simulations at each sample size, and used $\alpha = 0.05$ as a significance level.

### S4 Summary of published results

Table 1 of the main text summarizes results from empirical studies that estimate the heritability of individual taxa, for different host systems, populations and tissues. Below, we give some detailed information on each of the studies, and how we arrived at the numbers shown in Table 1. Where possible, we analyzed the provided heritability effect sizes and p-values, and use the same methodology to calculate the overall proportion of heritable microbes, in order to make results as comparable as possible. We note that the resulting estimates do not necessarily correspond to the results that are reported in the original study (for instance, some studies do not report the overall proportion of significant results, or use no or a different method to correct for multiple testing).

#### S4.1 O'Connor et al. (2014)

Host: Mice (gut microbiome)
Samples: 32 (8 outbred mice strains, 2 cages/strain, 4 animals/cage; divided over two diets. Only control diet is used for microbiome analyses)
Host genetic information: Lineage
Taxa: 43 taxa (genus level)
Transformation: Total Sum Scaling
Model: Heritabilities are estimated using inbred strain analysis, using relative abundances as the response variable, and no other covariates. Heritability is calculated as: $(0.5 * V_a)/(0.5 * V_a + V_e)$
Results: Heritability estimates range between 0.19 and 0.84 (average=0.47). We obtained these numbers by digitizing Table 1 in O'Connor et al. (2014). No significance or uncertainty measures are provided.

#### S4.2 Zhao et al. (2013)

Host: Chickens (gut microbiome)
Samples: 56 (consisting of males and females, from a low high and low body weight selection treatment)
Host genetic information: Pedigree
Taxa: 23 OTUs (from which 13 belong to Lactobacillus)
Transformation: Log-transformed and scaled
Model: Wombat software was used to obtain heritability estimates. No information on any other covariates is given, nor in how significance was assessed.
Results: None of the taxa is significantly heritable (note that no information on significance test is provided).

#### S4.3 Davenport et al. (2015)

Host: Humans (gut microbiome)
Samples: 93, 91 and 127 (for winter, summer, and both seasons combined, respectively)
Host genetic information: SNPs
Taxa: 16, 104 and 102 (for winter, summer, and both seasons combined, respectively. From genus to phylum level, including taxa that were present in at least 75% of the individuals, and excluding taxa that were highly correlated with a lower level or same level taxon ($r \geq 0.9$)).
Transformation: Quantile normalization
Model: Abundance data were collected for two seasons (summer and winter). For each season and per taxon, a quantile normalization was performed. For the analyses for both seaons combined, values were averaged for

individuals that were sampled in both seasons. Multiple regression with age, sex and date of collection on these normalized abundances was performed. Residuals were used to estimate the 'chip heritability' using GEMMA software based on ~200k SNPs. Results were considered significant if standard errors did not overlap with 0.
Results: 14/116 (12%), 10/104 (9.6%) and 13/102 (13%) of the taxa were heritable, for winter, summer and both seasons combined, respectively. From these heritable taxa, the average heritability was 0.35 (0.18-0.69), 0.37 (0.20-0.49) and 0.26 (0.19-0.37), for winter, summer and both seasons, respectively (calculated using provided Table S2 in Davenport et al. (2015)).

#### S4.4 Goodrich et al. (2014), data from Turnbaugh et al. (2009)

Host: Humans (gut microbiome)
Samples: 108 (23 DZ twin pairs, 31 MZ twin pairs).
Host genetic information: Twins
Taxa: 221 (149 OTUs, 72 higher-level taxonomic groups)
Transformation: Box-Cox transformation
Model: Same method as in S4.17. If two samples were collected from the same individual, randomly use one sample. Covariates: ancestry and the number of sequences per sample.
Results: We used Supplementary Table S2A from Goodrich et al. (2014), including all rows for Turnbaugh, using methods described in S4.17. 0% is heritable (0/221).

#### S4.5 Goodrich et al. (2014), data from Yatsunenko et al. (2012)

Host: Humans (gut microbiome)
Samples: 126 (34 DZ twin pairs, 29 MZ twin pairs).
Host genetic information: Twins
Taxa: 2933 (2664 OTUs, 269 higher-level taxonomic groups).
Transformation: Box-Cox transformation
Model: Same method as in S4.17, include only individuals aged >12. Covariates: age, the number of sequences per sample and sex.
Results: We used Supplementary Table S2A from Goodrich et al. (2014), including all rows for Yatsunenko, using methods described in S4.17. 0% is heritable (0/2933).

#### S4.6 Wright et al. (2021)

Host: Humans (vaginal microbiome)
Samples: 244 and 88 (European and African ancestry, respectively. Both MZ and DZ twin pairs)
Host genetic information: Twins
Taxa: 3 taxa (present in >20% of the samples and with a within-host proportion of >10%)
Transformation: Arcsine square root transformation on relative abundances
Model: OpenMx R-package was used to estimate heritabilities, for each self-identified ancestral group separately. Alpha level of 0.05 was used to assess significance.
Results: European ancestry: 1 taxon is significantly heritable (estimate: 0.35). African ancestry: no significant effects.

##### S4.7 Xie et al. (2016)

Host: Humans (gut microbiome)
Samples: 250 (35 MZ and 92 DZ twin pairs).
Host genetic information: Twins
Taxa: 109 genera (include when detected in >50% of the samples)
Transformation: Box-Cox transformation
Model: ACE model, implemented in OpenMx software, was fitted on the on Box-Cox transformed (1 as offset) values, correcting for sequencing amount, age and age started living apart. Obtain p-values by comparing ACE model with CE model, and correct for multiple testing according to Storey's FDR method.
Results: 11/109 of the genera showed a significant heritability (p-value < 0.1) (10%). Heritability estimates of individual taxa are not provided.

##### S4.8 Turpin et al. (2016)

Host: Humans (gut microbiome)
Samples: 270 (from 123 families with at least 2 individuals; this is a subset of the total sample size of 1,561 individuals).
Host genetic information: Pedigree
Taxa: 166 (excluding taxa with residuals Kurtosis ¿ 0.8, results in 249 taxa from phylum to genus level. Redundant taxa were then removed, leaving 166 taxa for the analyses)
Transformation: Inverse normal transformation
Model: Sequential Oligogenic Linkage Analysis Routines (SOLAR) was used to estimate heritabilities, adjusting for age and sex. Correct for correlations among taxa and multiple testing following Gao et al. (2010, 2008).
Results: 26/249 (10%) taxa have corrected p-value < 0.1. Note that Turpin et al. (2016) use a significance threshold of 0.05 (resulting in 20 significant taxa). From the 26 significant taxa (corrected $p < 0.1$), heritable ranges from 0.48 to 0.67 (average: 0.58).

##### S4.9 Sutherland et al. (2021)

Host: Switchgrass (rhizosphere)
Samples: 383 (3-10 clonally propagated individuals from 63 populations).
Host genetic information: SNPs
Taxa: 110 (families, >10% prevalence)
Transformation: Total Sum Scaling
Model: R-package *Sommer* was used to fit multivariate linear mixed models, using a kinship matrix based on the SNPs. Results was considered significant if standard errors did not overlap with zero.
Results: 21/110 families (19%) were significantly heritable, with estimates ranging from 0.106-0.539 (mean: 0.241).

##### S4.10 Wallace et al. (2019)

Host: Cattle (rumen microbiome)
Samples: 650 and 200 (from Holstein-Friesian and Nordic-Red breed, respectively).
Host genetic information: SNPs
Taxa: 512 (OTUs, across 11 prokaryotic orders, present in > 50% of the animals within each of the seven farms studied)

Transformation: Quantile normalization

Model: Genetics Complex Trait Analysis (GCTA) were used to calculate the variance in quantile normalized data that is explained by SNPs. Farm, dietary components and first five PCs of genetic relatedness matrix were used as covariates. Heritability confidence intervals were estimated based on software FIESTA, and p-values were corrected with the Benjamini-Hochberg approach (FDR<0.05).

Results: For Holstein-Friesian breed, 39/512 (7.6%) was heritable (ranges: 0.2-0.6). For Nordic-Red breed, 3/512 (0.59%) was heritable. Note that as the study does not provide p-values for each of the 512 taxa, we could not calculate the number of significant results when setting the FDR to 0.1, to make it more consistent with the other results.

##### S4.11 Gomez et al. (2017)

Host: Humans (oral microbiome)

Samples: 485 (205 MZ and 280 DZ twins between 5-11 years old)

Host genetic information: Twins

Taxa: 91 OTUs (present in at least 50% of the individuals)

Transformation: Log-transformed and scaled

Model: An ACE model was fitted using R-package mets to calculate heritabilities, controlling for sex and age.

Results: The supplementary Table S3 provides p-values for the variance explained by host genetics (A in ACE model). We applied the Benjamini Hochberg approach to correct p-values for multiple testing (FDR < 0.1). This results in 46% of the OTUs being significantly heritable (42/91 OTUs). Ranges: 0.13-0.65 (mean: 0.34). Note that Gomez et al. (2017) report 12 OTUs that were highly heritable with additive genetic control explaining >40% of the variation in abundance.

### S4.12 Si et al. (2017)

Host: Humans (vaginal microbiome)

Samples: 542 (222 MZ twins, 56 DZ twins, 102 family members, 80 related females and 82 unrelated females)

Host genetic information: Twins

Taxa: 369 (OTUs and higher taxonomic levels, included when present in > 50% and 15% of the individuals, respectively)

Transformation: Inverse normal transformation

Model: A linear regression was fitted on the inverse normally transformed abundances, correcting for age, menopause, hormone therapy, bacterial vaginosis, HPV, BMI, waist-hip ratio, OTU counts per sample and sequence run. Sequantial Oligogenic Linkage Analysis Routine (SOLAR) was then used to estimate heritabilities. The study confirms heritability estimates using OpenMx software.

Results: 85/369 of the taxa is significantly heritable (23%). We calculated this using Table S1a (Si et al., 2017), using the provided corrected p-values (FDR < 0.1). From these taxa, the average heritability is 0.27 (ranges: 0.13-0.49).

##### S4.13 Org et al. (2015)

Host: Mice (gut microbiome)

Samples: 592 samples (from 113 inbred strains, 7 samples were omitted from further analysis)

Host genetic information: SNPs

Taxa: 43 taxa (including only those present in >75% of the samples, from different taxonomic levels)
Transformation: Total Sum Scaling
Model: EMMAX software was used to estimate heritabilities, based on a mixed effects model with relative abundances as response variable. SNP-data were used to determine genetic relatedness, and no other covariates were included
Results: Combining both males and females, heritability estimates range between 0.26 and 0.77 (average=0.51) (numbers obtained from Table S3 in Org et al. (2015)). No significance or uncertainty measures are provided.

#### S4.14 Deng et al. (2021)

Host: Sorghum (rhizosphere)
Samples: 600 (200 genotypes, 3 blocks)
Host genetic information: Lineage
Taxa: 1189 OTUs (representing 29 phyla; removing low abundance OTUs with <3 reads in at least 20 % of the samples)
Transformation: Cumulative Sum Scaling
Model: Linear mixed effects model (R-package *Sommer*) was fitted, including random effect of replicate nested within block. The study accounts for spatial trends in the field, by excluding this source of variance for heritability calculations. P-values were calculated by permuation analyses, randomly shuffling OTU abundances 1000 times. In addition, Deng et al. (2021) use SNP data to estimate heritabilities. These are lower than the calculated broad-sense heritabilities, but p-values or standard errors are not provided, and are therefore not included in our Table 1.
Results: The p-values obtained by permutation analyses are not provided. Instead, Table S4 in Deng et al. (2021) reports standard errors of the heritability estimates. For 443/1189 OTU's (37%), standard errors do not overlap with 0. From these OTUs, the average heritability is 0.22, ranging between 0.11 and 0.66.

#### S4.15 Lim et al. (2017)

Host: Humans (gut microbiome)
Samples: 655 (153 MZ twin pairs, 37 DZ twin pairs, and 275 parents and siblings).
Host genetic information: Twins
Taxa: 85 (excluding taxa with <0.1% of mean relative abundance)
Transformation: Inverse normal transformation
Model: Sequential Oligogenic Linkage Analysis Routines (SOLAR) was used to estimate heritabilities, with the following covariates: age, sex and disease status (Metabolic syndrome). P-values were corrected using Benjamini-Hochberg approach.
Results: 52/85 taxa (61%) have a non-zero heritability (q-value < 0.1) (calculated using Supplementary Table S3 in Lim et al. (2017)). Lim et al. (2017) report taxa with a corrected p-value < 0.05. For consistency with the other studies discussed here, we use a FDR of 0.1. However, we note that when using a threshold of 0.05, we obtain 50 significant taxa, so results are very similar. From these significant taxa, heritable ranges from 0.13 to 0.46 (average: 0.24).

#### S4.16 Ishida et al. (2020)

Host: Humans (gut microbiome)
Samples: 1068

Host genetic information: SNPs
Taxa: 21 genera (found in >95% of the samples)
Transformation: Box-Cox transformation
Model: Following methods as described in Davenport et al. (2015). Box-Cox transformed relative abundances were regressed on the sex variable, and residuals were implemented in GEMMA software to calculate the 'chip heritability', i.e. the variance explained by genome-wide SNPs. Significance was based on error bars not intersecting with zero.
Results: 6/21 of the genera are significantly heritable (29%). Estimates of significant genera range between 0.33 and 0.54 (average: 0.40).

#### S4.17 Goodrich et al. (2014)

Host: Humans (gut microbiome)
Samples: 1081 (Total of 977 individuals from TwinUK Dataset. 171 MZ and 245 DZ twin pairs; 2 from twin pairs with unknown zygosity; 143 samples from one twin).
Host genetic information: Twins
Taxa: 909 (768 OTUs that were present in >50% of the samples, and higher-level taxonomic groups)
Transformation: Box-Cox transformation
Model: Multiple regression on Box-Cox transformed (offset of 1) trait abundances, with the following covariates: gender, age, number of OTU counts per sample, technician identity, sequencing run and shipment batch. Use the residuals of this model to fit an ACE model (using software OpenMx), to decompose the total variance into the contribution of genetic effects (A), common environment (C) and unique environment (E). P-values were obtained by permuting MZ and DZ labels 10,000 times, and calculate heritability for each permuted dataset. Calculate the proportion at which the permuted heritability met or exceeded the observed heritability. Correct for multiple testing using Benjamini-Hochberg approach, using a FDR of 0.1
Results: We used Supplementary Table S2A to calculate the proportion of significant microbes. Include all rows for TwinUK, correct p-values using Benjamini-Hochberg approach, using an FDR < 0.1. 1.1% is heritable (10/909). Including only significantly heritable microbes, the average heritability is 0.29 (range: 0.09 - 0.39).

#### S4.18 Kurilshikov et al. (2021)

Host: Humans (gut microbiome)
Samples: 1176 (169 MZ and 419 DZ twin pairs; from TwinsUK dataset)
Host genetic information: Twins
Taxa: 159 (different taxonomic levels; we note that the provided supplementary table reports estimates for 209 taxa)
Transformation: Inverse rank-sum transformation
Model: An ACE-model was fitted using the OpenMx package, refering to Goodrich et al. (2014) for more details. We used the provided p-values to correct for multiple testing (FDR < 0.1).
Results: 11/209 (5%) of the taxa is heritable. The average non-zero heritability is 0.31 (ranges: 0.27-0.40).

#### S4.19 Bergamaschi et al. (2020)

Host: Pigs (gut microbiome)

Samples: 1205, 1295 and 1283 (28 families, animals were sampled during three different time points; 1039 animals were sampled at all three time points).
Host genetic information: Pedigree
Taxa: 1678 OTUs
Transformation: Total sum scaling
Model: OTU counts (transformation not mentioned, assuming relative abundances are used, i.e. total sum scaling) were used in a sire model. Sex, sire, dam and contemporary group were included as fixed effects; sire and pen were included as random effects. Heritability calculated as: $(4V_{sire})/(V_{sire} + V_{pen} + V_E)$. Significance level is not provided; we assume that heritabilies are considered significant when standard errors do not overlap with 0.
Results: Heritability varied over the 3 time points, increasing over lifespan. Out of the 1678 OTUs, 170, 261, 366 were heritable (10%, 16% and 22%), for the three time points, respectively. Ranges are: 0.025-0.14, 0.029-0.29, and 0.025-0.545, with averages 0.056, 0.078 and 0.099, for the three time points, respectively.

#### S4.20 Walters et al. (2018)

Host: Maize (rhizosphere)
Samples: 4866 and 45 for a field study in 2010 and 2015, respectively (2010: 27 parental genotypes, 2015: 5 parental genotypes).
Host genetic information: Lineage
Taxa: 792 and 2557 OTUs (2010 and 2015 field study, respectively. Present in >80% of the samples)
Transformation: Log-transformation on relative abundances
Model: Mixed effects models (using lme4) were fitted on the log-transformed relative abundances. Sampling depth was included as fixed effect, and week, location and inbred line were included as random effects. Broad-sense heritability was calculated as the proportion of variance explained by inbred line. To obtain p-values, a permutation analysis (permuting abundances across all samples, 5000 times) was performed. Heritabilities were considered significant if $p \leq 0.001$.
Results: For the 2010 field study: 143/792=18% OTUs are significantly heritable (with $p<0.001$). Estimates of significant taxa range between 0.15-0.23, with an average of 0.17 (calculated using Dataset S4 in Walters et al. (2018)). For the 2015 field study: 5/2557 (0.2%) of the OTUs are heritable ($p < 0.001$), with estimates ranging between 0.41 and 0.53 (mean: 0.45) (calculated using Dataset S4). Note that only 200 the OTUs with highest heritability are provided in Dataset S4, so we could not correct p-values using Benjamini Hochberg approach, and assess the significance using the same approach as done in other studies.

#### S4.21 Goodrich et al. (2016)

Host: Humans (gut microbiome)
Samples: 3261 samples (2731 different individuals including 489 DZ twin pairs and 637 MZ twin pairs. 530 individuals with at least two samples).
Host genetic information: Twins
Taxa: 945 taxa (from which 782 OTUs; include taxa that are present in >50% of the samples)
Transformation: Box-Cox transformation
Model: Use Box-Cox transformed (offset of 1) relative abundances in multiple regression, with the following covariates: number of 16s rRNA sequences per sample, age, gender, shipment date, collection method, and technician identity. Residuals were used in ACE model, implemented in OpenMx software. Significance of host genetics was determined by comparing with CE model (without host genetics), based on a likelihood ratio test. For samples

from the same individual, only first measurement was included. Use Benjamini-Hochberg approach to correct for multiple testing.
Results: We used the adjusted p-values provided in Table S1.52 to calculate the proportion of heritable taxa. 52 taxa have a corrected p-value $< 0.1$, which is 5.5% of all tested taxa (52/945). Heritability estimates (including significant taxa) range between 0.1-0.42, with an average of 0.3. Note: in the original paper, Goodrich et al. (2016) focus more on effect sizes, discussing heritability estimates $> 0.2$.

#### S4.22 Gacesa et al. (2020)

Host: Humans (gut microbiome)
Samples: 4745 (in 2756 parent-child pairs, 530 sibling pairs and 815 pairs with second degree or more distant relationship)
Host genetic information: Pedigree
Taxa: 242 (different taxonomic levels, including taxa present in at least 5% of the samples and with a mean relative abundance $> 0.01\%$)
Transformation: Centered log-ratio transformation
Model: Broad-sense heritability is estimated, adjusting for age, sex, BMI, read depth and stool frequency. P-values were corrected using Benjamini-Hochberg approach, and considered significant when FDR $< 0.1$.
Results: 31 of the 241 taxa are significantly heritable (12.9%). From these taxa, heritability estimates range between 0.15 and 0.30 (average=0.20).

#### S4.23 Grieneisen et al. (2021)

Host: Baboons (gut microbiome)
Samples: 16,234 (longitudinal data for 14 years on total of 585 baboons; each individual on average 28 samples).
Host genetic information: Pedigree
Taxa: 283 (139 ASVs and 144 higher-level taxonomic groups found in >50% of the samples, including only the relative abundances as phenotypic traits (single-taxon phenotypes). The study includes a total of 1034 phenotypes, additionally including seven community phenotypes and 744 absence/presence phenotypes found in 10-90% of the samples).
Transformation: Total Sum Scaling (Centered Log-Ratio transformation and Phylogenetic Balances transformation are also performed, but not included here. We note that results were very similar.)
Model: Fit animal model using the R-package ASReml. Random effects of individual, maternal identity, DNA extraction plate, and additive genetics effect. Fixed effects of diet (13 principal components), age, date, month, rainfall, hydrological year, social group membership, group size, post-PCR DNA concentration and read count. Calculate heritability after correcting for fixed effects. Significance assessed by comparing to model without host genetic effects, based on a log-likelihood ratio test. Correct for multiple testing using Benjamini-Hochberg approach, using a FDR of 0.1.
Results: 93% of the single-taxon phenotypes (273/283) is heritable. Using a FDR of 0.01 (instead of 0.1) gives similar results. Average significant heritability is 0.068, ranging between 0.008 and 0.21.

#### S5 Data transformations

Using relative microbial abundances (where the sum in each sample is set to 1) can result in an overestimation of the proportion of heritable microbes (Fig. 2 in the main text and Fig. S1). Instead of using relative abundances, one could also normalize data based on a chosen normalization factor. In addition, expressing abundances as log ratios compared to a reference, transforms data to real space, making it suitable for regression analyses. Below we apply an Additive Log-Ratio (ALR) transformation to explore its consequences, making use of simulated datasets. Here, the abundance of each taxon $i$ ($x_i$) is divided by the abundance of a reference taxon, followed by taking the logarithm: $\mathrm{ALR}(x_i) = \log(x_j/x_{\mathrm{ref}})$ (Greenacre et al., 2021).

When there truly is a reference taxon that has a constant abundance across samples (hence, $V_{P\mathrm{ref}} = 0$), applying an ALR transformation can succesfully resolve the issue of high false discovery rates (Fig. S2), resuting in accurate estimates of the overall proportion of heritable microbes. When the set reference varies slightly in abundance across hosts (10% of the variance compared to the other taxa), it is still possible to control for high false discovery rates. However, this comes at the cost of some statistical power at lower sample sizes resulting in an underestimation of the proportion of heritable microbes. This decrease in statistical power becomes more pronounced when the chosen reference taxon has the same variance in abundance as the other taxa (and is thus not a true reference taxon, although not under host genetic control).

Choosing a reference taxon that, in reality, is heritable itself, is more problematic. Dividing each microbial abundance by a taxon's abundance that varies across host genotypes, creates a spurious genotypic signal in each taxon. Indeed, except when sample sizes are very low (and thus low statistical power), this leads to the wrong conclusion that all microbes are heritable, even if the reference taxon varies only slightly in abundance across hosts.

This illustrates the importance of choosing appropriate normalization factors. When there truly is a reference taxon, one could succesfully resolve the problem of high false discovery rates. On the other hand, if the chosen reference taxon is under some host genetic control itself ($h^2 > 0$), the problem could become even worse.

Other transformations divide each taxon count by some other factor, e.g. by the geometric mean abundance within each sample (Centered Log-Ratio transformation), implicitely assuming that the geometric mean abundance is constant across samples. It is beyond the scope of our paper to test all transformations and statistical methods, but there is a large body of literature on this in the context of differential abundance analyses (Hawinkel et al., 2019; Lin and Peddada, 2020). While some methods may be more suitable than others, as the appropriate reference values will depend on the biology, there unlikely is a one-size-fits-all recommended transformation. Moreover, as has been shown with many of these methods, it remains challenging to control for high false discovery rates (Hawinkel et al., 2019; Lin and Peddada, 2020).

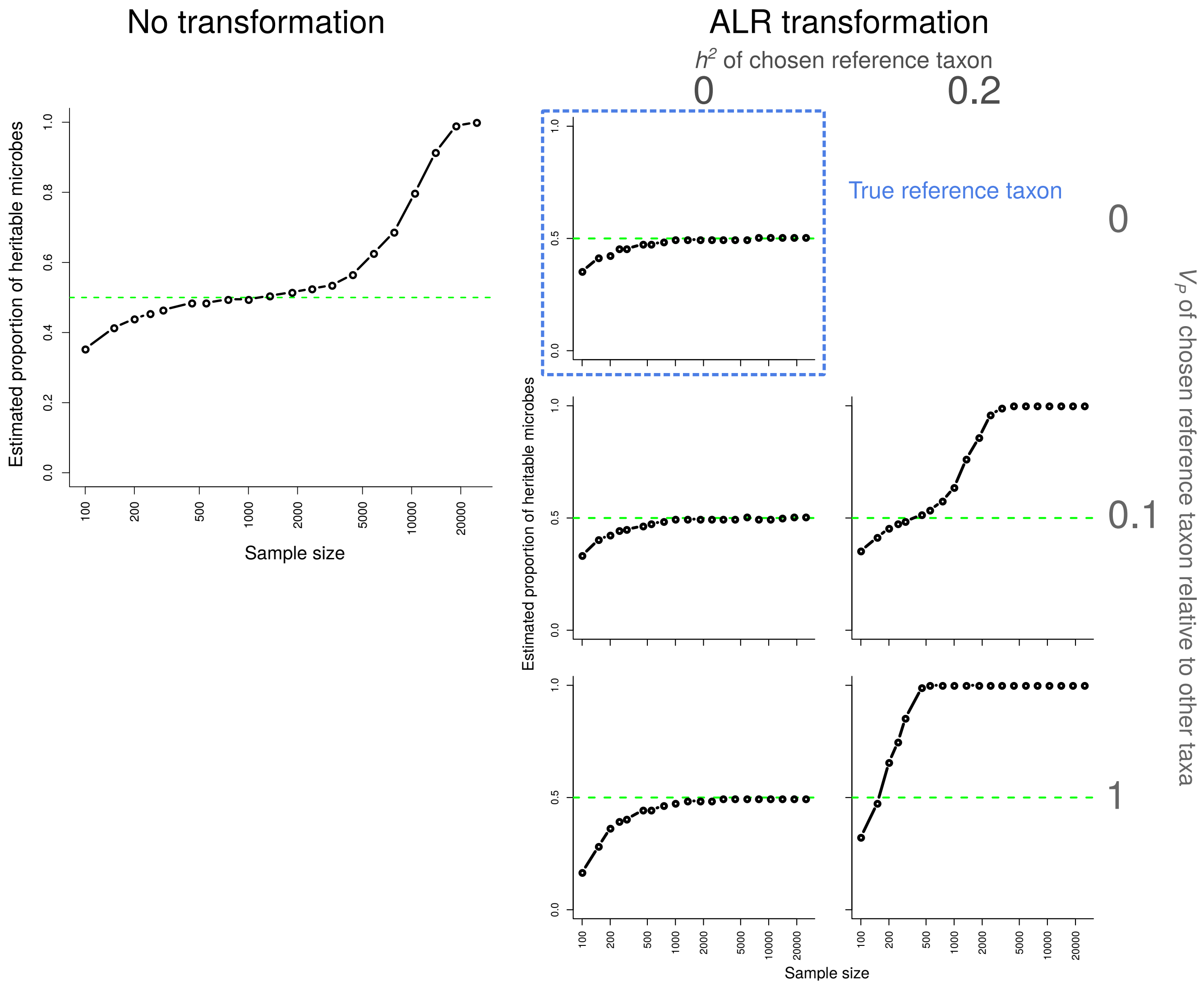

Figure S2: Estimated proportion of heritable microbes (y-axes) as a function of sample size (x-axes) when the true proportion of heritable microbes is set at 0.5 (green lines). Left: no transformation is applied and analyses are thus performed on the simulated relative abundances. Right: results when applying an ALR transformation, expressing abundances as log ratios compared to a chosen reference taxon. Here we vary properties of the reference taxon (i.e. its relative variance across hosts [rows], and its heritability [columns]). Note that only a taxon with a variance of 0, truly is a reference taxon, i.e. that is stable across hosts. We simulated communities consisting of 100 microbes, each with an average absolute abundance of 1, following the procedure described in Section S2.1. Dots show median values of 100 simulations.

#### References

M. Bergamaschi, C. Maltecca, C. Schillebeeckx, N. P. McNulty, C. Schwab, C. Shull, J. Fix, and F. Tiezzi. Heritability and genome-wide association of swine gut microbiome features with growth and fatness parameters. *Scientific reports*, 10(1):1–12, 2020.

E. R. Davenport, D. A. Cusanovich, K. Michelini, L. B. Barreiro, C. Ober, and Y. Gilad. Genome-wide association studies of the human gut microbiota. *PloS one*, 10(11):e0140301, 2015.

S. Deng, D. F. Caddell, G. Xu, L. Dahlen, L. Washington, J. Yang, and D. Coleman-Derr. Genome wide association study reveals plant loci controlling heritability of the rhizosphere microbiome. *The ISME Journal*, pages 1–14, 2021.

R. Gacesa, A. Kurilshikov, A. V. Vila, T. Sinha, M. Klaassen, L. Bolte, S. Andreu-Sánchez, L. Chen, V. Collij, S. Hu, J. Dekens, V. Lenters, J. Björk, J. Swarte, M. Swertz, B. Jansen, J. Gelderloos-Arends, L. cohort Study, M. Hofker, R. Vermeulen, S. Sanna, H. Harmsen, C. Wijmenga, J. Fu, A. Zhernakova, and R. Weersma. The Dutch Microbiome Project defines factors that shape the healthy gut microbiome. *bioRxiv*, page 2020.11.27.401125, nov 2020. doi: 10.1101/2020.11.27.401125. URL `https://www.biorxiv.org/content/10.1101/2020.11.27.401125v1https://www.biorxiv.org/content/10.1101/2020.11.27.401125v1.abstract`.

X. Gao, J. Starmer, and E. R. Martin. A multiple testing correction method for genetic association studies using correlated single nucleotide polymorphisms. *Genetic Epidemiology: The Official Publication of the International Genetic Epidemiology Society*, 32(4):361–369, 2008.

X. Gao, L. C. Becker, D. M. Becker, J. D. Starmer, and M. A. Province. Avoiding the high Bonferroni penalty in genome-wide association studies. *Genetic Epidemiology: The Official Publication of the International Genetic Epidemiology Society*, 34(1):100–105, 2010.

A. Gomez, J. L. Espinoza, D. M. Harkins, P. Leong, R. Saffery, M. Bockmann, M. Torralba, C. Kuelbs, R. Kodukula, J. Inman, and Others. Host genetic control of the oral microbiome in health and disease. *Cell host \& microbe*, 22(3):269–278, 2017.

J. K. Goodrich, J. L. Waters, A. C. Poole, J. L. Sutter, O. Koren, R. Blekhman, M. Beaumont, W. Van Treuren, R. Knight, J. T. Bell, and Others. Human genetics shape the gut microbiome. *Cell*, 159(4):789–799, 2014.

J. K. Goodrich, E. R. Davenport, M. Beaumont, M. A. Jackson, R. Knight, C. Ober, T. D. Spector, J. T. Bell, A. G. Clark, and R. E. Ley. Genetic determinants of the gut microbiome in UK twins. *Cell host & microbe*, 19 (5):731–743, 2016.

P. Green and C. J. MacLeod. simr: an R package for power analysis of generalised linear mixed models by simulation. *Methods in Ecology and Evolution*, 7(4):493–498, 2016. doi: 10.1111/2041-210X.12504. URL `https://cran.r-project.org/package=simr`.

M. Greenacre, M. Martinez-Álvaro, and A. Blasco. Compositional Data Analysis of Microbiome and Any-Omics Datasets: A Validation of the Additive Logratio Transformation. *Frontiers in Microbiology*, page 2625, 2021.

L. Grieneisen, M. Dasari, T. J. Gould, J. R. Björk, J.-C. Grenier, V. Yotova, D. Jansen, N. Gottel, J. B. Gordon, N. H. Learn, and Others. Gut microbiome heritability is nearly universal but environmentally contingent. *Science*, 373(6551):181–186, 2021.

X. A. Harrison, L. Donaldson, M. E. Correa-Cano, J. Evans, D. N. Fisher, C. E. D. Goodwin, B. S. Robinson, D. J. Hodgson, and R. Inger. A brief introduction to mixed effects modelling and multi-model inference in ecology. *PeerJ*, 6:e4794, 2018.

S. Hawinkel, F. Mattiello, L. Bijnens, and O. Thas. A broken promise: microbiome differential abundance methods do not control the false discovery rate. *Briefings in bioinformatics*, 20(1):210–221, 2019.

J. Hayya, D. Armstrong, and N. Gressis. A note on the ratio of two normally distributed variables. *Management Science*, 21(11):1338–1341, 1975.

S. Ishida, K. Kato, M. Tanaka, T. Odamaki, R. Kubo, E. Mitsuyama, J.-z. Xiao, R. Yamaguchi, S. Uematsu, S. Imoto, and Others. Genome-wide association studies and heritability analysis reveal the involvement of host genetics in the Japanese gut microbiota. *Communications biology*, 3(1):1–10, 2020.

A. Kurilshikov, C. Medina-Gomez, R. Bacigalupe, D. Radjabzadeh, J. Wang, A. Demirkan, C. I. Le Roy, J. A. R. Garay, C. T. Finnicum, X. Liu, et al. Large-scale association analyses identify host factors influencing human gut microbiome composition. *Nature Genetics*, 53(2):156–165, 2021.

M. Y. Lim, H. J. You, H. S. Yoon, B. Kwon, J. Y. Lee, S. Lee, Y.-M. Song, K. Lee, J. Sung, and G. Ko. The effect of heritability and host genetics on the gut microbiota and metabolic syndrome. *Gut*, 66(6):1031–1038, 2017.

H. Lin and S. D. Peddada. Analysis of microbial compositions: a review of normalization and differential abundance analysis. *NPJ biofilms and microbiomes*, 6(1):1–13, 2020.

J. Oberpriller, M. de Souza Leite, and M. Pichler. Fixed or random? On the reliability of mixed-effect models for a small number of levels in grouping variables. *bioRxiv*, 2021.

A. O'Connor, P. M. Quizon, J. E. Albright, F. T. Lin, and B. J. Bennett. Responsiveness of cardiometabolic-related microbiota to diet is influenced by host genetics. *Mammalian Genome*, 25(11):583–599, 2014.

E. Org, B. W. Parks, J. W. J. Joo, B. Emert, W. Schwartzman, E. Y. Kang, M. Mehrabian, C. Pan, R. Knight, R. Gunsalus, and Others. Genetic and environmental control of host-gut microbiota interactions. *Genome research*, 25(10):1558–1569, 2015.

J. Si, H. J. You, J. Yu, J. Sung, and G. Ko. Prevotella as a hub for vaginal microbiota under the influence of host genetics and their association with obesity. *Cell host & microbe*, 21(1):97–105, 2017.

J. Sutherland, T. Bell, R. V. Trexler, J. Carlson, and J. Lasky. Host genomic influence on bacterial composition in the switchgrass rhizosphere. *bioRxiv*, 2021.

P. J. Turnbaugh, M. Hamady, T. Yatsunenko, B. L. Cantarel, A. Duncan, R. E. Ley, M. L. Sogin, W. J. Jones, B. A. Roe, J. P. Affourtit, and Others. A core gut microbiome in obese and lean twins. *nature*, 457(7228):480–484, 2009.

W. Turpin, O. Espin-Garcia, W. Xu, M. S. Silverberg, D. Kevans, M. I. Smith, D. S. Guttman, A. Griffiths, R. Panaccione, A. Otley, and Others. Association of host genome with intestinal microbial composition in a large healthy cohort. *Nature genetics*, 48(11):1413–1417, 2016.

R. J. Wallace, G. Sasson, P. C. Garnsworthy, I. Tapio, E. Gregson, P. Bani, P. Huhtanen, A. R. Bayat, F. Strozzi, F. Biscarini, and Others. A heritable subset of the core rumen microbiome dictates dairy cow productivity and emissions. *Science advances*, 5(7):eaav8391, 2019.

W. A. Walters, Z. Jin, N. Youngblut, J. G. Wallace, J. Sutter, W. Zhang, A. González-Peña, J. Peiffer, O. Koren, Q. Shi, and Others. Large-scale replicated field study of maize rhizosphere identifies heritable microbes. *Proceedings of the National Academy of Sciences*, 115(28):7368–7373, 2018.

M. L. Wright, J. M. Fettweis, L. J. Eaves, J. L. Silberg, M. C. Neale, M. G. Serrano, N. R. Jimenez, E. Prom-Wormley, P. H. Girerd, J. F. Borzelleca, et al. Vaginal microbiome lactobacillus crispatus is heritable among european american women. *Communications Biology*, 4(1):1–6, 2021.

H. Xie, R. Guo, H. Zhong, Q. Feng, Z. Lan, B. Qin, K. J. Ward, M. A. Jackson, Y. Xia, X. Chen, and Others. Shotgun metagenomics of 250 adult twins reveals genetic and environmental impacts on the gut microbiome. *Cell systems*, 3(6):572–584, 2016.

T. Yatsunenko, F. E. Rey, M. J. Manary, I. Trehan, M. G. Dominguez-Bello, M. Contreras, M. Magris, G. Hidalgo, R. N. Baldassano, A. P. Anokhin, and Others. Human gut microbiome viewed across age and geography. *nature*, 486(7402):222–227, 2012.

L. Zhao, G. Wang, P. Siegel, C. He, H. Wang, W. Zhao, Z. Zhai, F. Tian, J. Zhao, H. Zhang, and Others. Quantitative genetic background of the host influences gut microbiomes in chickens. *Scientific reports*, 3:1163, 2013.
